# A gut-derived hormone switches dietary preference after mating in *Drosophila*

**DOI:** 10.1101/2022.03.02.482605

**Authors:** Alina Malita, Olga Kubrak, Takashi Koyama, Nadja Ahrentløv, Kenneth V. Halberg, Michael J. Texada, Stanislav Nagy, Kim Rewitz

## Abstract

Animals must adapt their dietary choices to meet their nutritional needs. How these needs are detected and translated into nutrient-specific appetites that drive food-choice behaviors is poorly defined. Here, we show that the enteroendocrine cells (EECs) of the adult female *Drosophila* midgut sense nutrients and in response release neuropeptide F (NPF), an ortholog of mammalian NPY-family gut-brain hormones. Gut-derived NPF acts via effects on glucagon-like adipokinetic hormone (AKH) signaling to induce sugar satiety and to drive hunger for protein-rich food, and on adipose tissue to promote storage of ingested nutrients. Suppression of gut NPF leads to overconsumption of dietary sugar while decreasing intake of protein-rich yeast. Furthermore, we show a female-specific function of gut-derived NPF in the suppression of AKH signaling after mating. This induces a dietary switch that promotes preference for protein-containing food to support reproduction. Together, our findings suggest that the gut NPF-AKH axis regulates appetite that drives specific food choices to ensure homeostatic consumption of nutrients, providing insight into the hormonal mechanisms that underlie nutrient-specific hungers.

## Introduction

Animals must be able to select the specific nutrients they need to consume. Food selection is governed by appetites for specific nutrients to ensure adequate ingestion of macronutrients needed to maintain nutritional homeostasis and optimal fitness ^1,2^. This has given rise to the hypothesis that organisms can feel specific hungers or appetites for the type of nutrients they need ^3,4^. Nutrient-specific appetite is the drive to eat food containing specific nutrients, and this behavior has been demonstrated in many organisms, including humans ^3-6^. Such homeostatic nutrient consumption requires sensors that detect the internal nutritional state and mechanisms that translate this information into changes in feeding decisions. Food consumption is controlled by nutritional signals from the periphery, such as the adipokine leptin and a variety of gut hormones, that act together with circulating nutrients on the brain ^7^. However, the hormones and mechanisms that govern nutrient-specific appetites that drive appropriate food choices to restore homeostasis are poorly defined.

The fruit fly *Drosophila*, like mammals, regulates feeding behaviors according to internal state ^1,6,8,9^. Because of the metabolic demands of egg production, food preferences are sex-specific and strongly dependent on mating status ^6^. While nutrient-replete flies prefer sugar-rich foods, protein-deprived animals shift their preference towards foods rich in yeast, which is their main source of dietary protein. Mated female flies need protein to support egg production, and they therefore more strongly prefer to consume foods rich in yeast than both males and virgin females. This switch in preference towards protein-rich food is driven by male-produced Sex Peptide (SP), which is transferred to the female reproductive tract in seminal fluid during mating ^6^ and acts indirectly on the midgut ^6,10^. Thus, internal energy state and mating status create drives that direct food-seeking behavior and consumption of specific nutrients.

Adjustment of feeding behaviors is mediated by satiety and hunger signals from peripheral organs, such as the gut. The gut is one of the largest endocrine organs, releasing a large number of different hormones from specialized enteroendocrine cells (EECs) in both flies and mammals ^11,12^. Gut-to-brain signaling conveys important information about the nutritional nature of the contents of the intestine, and intestinal nutrient-sensing and signaling play key roles in regulating food intake ^13,14^. For example, in the nutrient-deficient state, orexigenic or hunger signals from the mammalian gut such as ghrelin drive appetite to promote food consumption. Conversely, in response to food consumption, the mammalian gut releases glucagon-like peptide 1 (GLP-1), which acts as a satiety signal that reduces food intake. Satiety signals prevent excess nutrient intake, which can lead to the development of obesity and associated metabolic disorders, and GLP-1 therapy is effective in reducing body weight by lowering appetite ^15^. The fly gut is structurally similar to the mammalian gastrointestinal tract, and many gut-derived hormones are evolutionarily conserved ^16,17^, making *Drosophila* an attractive model for unraveling the signals by which the gut controls feeding decisions and sex differences in feeding behavior.

While significant progress has been made in understanding gut hormones that control metabolism ^18-20^, much less is known about how the gut communicates the presence of specific nutrients to adjust food choice, and gut signals that regulate appetite towards specific nutrients have not been described. In particular, excessive sugar consumption is a main cause of obesity, yet the mechanisms by which the gut senses sugar and regulates carbohydrate satiety are poorly defined. Here, we show that a population of EECs in the adult female *Drosophila* gut sense sugar via a mechanism requiring Sugar transporter 2 (Sut2) and release Neuropeptide F (NPF), an ortholog of mammalian NPY hormones. We find that the gut-derived NPF acts via several routes of tissue crosstalk on the Allatostatin C (AstC)-adipokinetic hormone (AKH) axis to suppress sugar intake and drive preference towards protein-rich food after mating, suggesting that NPF controls sex difference in food intake required for reproductive processes. Our findings suggest that NPF and AKH hormonal signaling drives food choices that ensure homeostatic consumption by promoting sugar satiety and protein-specific hunger.

## Results

### Midgut NPF suppresses sugar intake and energy breakdown in females

To identify gut-derived hormones and nutrient-sensing mechanisms that regulate feeding, we performed an *in vivo* RNA-interference screen of secreted factors and receptors in adult *Drosophila*. We focused on the EECs, which produce a variety of factors that have key roles in the coordination of food intake and metabolism ^13,18-20^. We examined the effect of adult-restricted, EEC-specific hormone knockdown on male and female sugar-water feeding behavior using the FLIC (Fly Liquid-food Interaction Counter) system ^21^ (Fig. 1A). This apparatus allows automated monitoring of *Drosophila* feeding behaviors by detecting when flies are in physical contact with the food. We used the driver *voilà-GAL4* to target the RNAi effect to the EECs, in combination with temperature-sensitive (TS) *Tub-GAL80* (*GAL80*^*TS*^), together referred to as *EEC*> hereafter, which allowed us to induce gene silencing only in the adult stage ^18,20^. Among our hits was the peptide NPF, knockdown of which increased the time mated females spent in contact with sugar-only food (Fig. 1B) while decreasing males’ food-interaction time (Fig. S1A). To rule out contributions of the UAS transgene itself to the phenotype, we crossed the *UAS-NPF-RNAi* (*NPFi*) line to the *w*^*1118*^ background; this genotype showed results similar to those seen with the driver control (Fig. S1B). This suggests that lack of NPF production in the EECs of adult females enhances their interest in or motivation to feed on sugar. Next, we measured short-term (30-minute) food intake using a dye-consumption assay. In this case, we used standard adult fly food ^22^ containing both sugar (9%) and yeast (8%) along with a second RNAi construct targeting NPF to confirm the specificity of the observed effects. To prime animals for feeding, we preconditioned them by fasting them 15 hours and performed all food intake experiments during time when animals have their morning meal (one hour after lights on; 12:12 hour dark cycle). Females with EEC suppression of *NPF* consumed significantly more sugar+yeast food than controls (Fig. 1C and S1C). To measure sugar consumption, we used the capillary feeder (CAFÉ) assay ^23^ to quantify food intake over 6 hours. Females with adult-restricted EEC knockdown of *NPF* consumed significantly more sugar over the 6-hour period than controls (Fig. 1D and S1D). In males, loss of *NPF* in the EECs led to reduced food consumption over an even longer period of 24 hours (Fig. S1E), consistent with the observed decrease in their food-interaction time (Fig. S1A). Together these results suggest that gut-derived NPF suppresses postprandial sugar intake in females.

**Figure 1.**
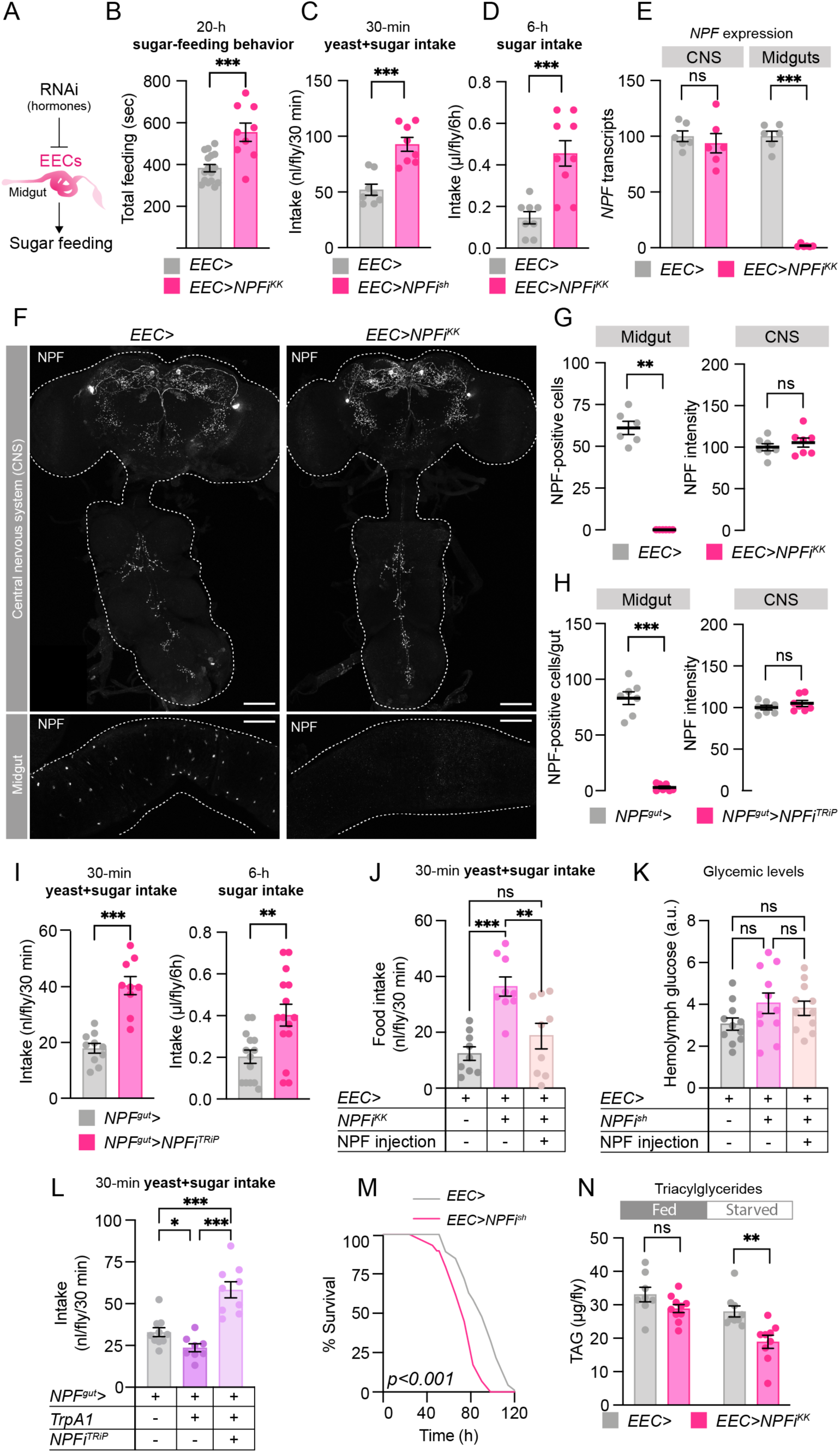
Gut-derived NPF regulates sugar intake and metabolism in mated females. (A) Sugar-feeding behavior was measured in animals expressing RNAi-mediated knockdown of hormones in the enteroendocrine cells (EECs) of the midgut. (B) Total time of mated females spent feeding on 10%-sugar food over 20 hours, measured using FLIC. Student’s t-test; n=16 *EEC*>, n=9 *EEC*>*NPFi*^*KK*^. (C and D) Consumption of 10% sugar over 6 hours measured by CAFÉ assay (C) and of sugar+yeast food (9% sugar and 8% yeast) over 30 minutes determined by dye assay (D) of mated female flies after 15-hour starvation. C, Student’s t-test; n=8 *EEC*>, n=9 *EEC*>*NPFi*^*KK*^. D, Student’s t-test; n=8 *EEC*> and *EEC*>*NPFi*^*sh*^. (E) The conditional *EEC*> driver used to knock *NPF* down in the EECs efficiently reduced *NPF* transcripts in the midgut without affecting expression in the central nervous system (CNS; brain and ventral nerve cord, VNC) of mated females. One-way Kruskal-Wallis ANOVA; n=6 independent biological replicates from tissues pooled from 6 animals for each condition. (F and G) NPF immunostaining of CNS and midgut tissue from mated females with knockdown of *NPF* reveals that *EEC*>-driven RNAi eliminates detectable NPF in the midgut but does not reduce CNS levels, quantified in G. Mann-Whitney U test; n=7 CNS, n=6 midguts. Scale bars, 50 µm. (H) Immunostaining shows that *NPF* knockdown using the *NPF*> driver with pan-neuronal *GAL80* (*R57C10-GAL80; NPF*>: *NPF*^*gut*^>) reduces NPF specifically in the midgut EECs and not in the CNS. Quantification of images represented in Fig. S1I. Mann-Whitney U test; n=7 tissues each. (I) Consumption of sugar+yeast food over 30 minutes measured by dye assay and of 10% sugar over 6 hours measured by CAFÉ assay of 15-hour-starved flies. Student’s t-test; left n=10 *NPF*^*gut*^>, n=9 *NPF*^*gut*^>*NPFi*^*TRiP*^, right n=14 *NPF*^*gut*^>, n=15 *NPF*^*gut*^>*NPFi*^*TRiP*^. (J and K) Injection of NPF peptide into the hemolymph of 15-hour-starved mated females restores normal feeding without affecting hemolymph glucose levels. (J) Consumption of sugar+yeast food over 30 minutes in 15-hour-starved mated females. One-way ANOVA with Tukey’s test; n=9 each for *EEC*>, *EEC*>*NPFi*^*KK*^, *EEC*>*NPF*^*KK*^*+*NPF peptide. (K) One-way ANOVA with Tukey’s multiple comparisons test; n=11 each. (L) Activation of NPF^+^ EECs by expressing the heat-sensitive TrpA1 channel decreases 30-minute intake of sugar+yeast food (9% sugar and 8% yeast) measured by dye assay in an NPF-dependent manner in 15-hour-starved mated females. One-way ANOVA with Tukey’s multiple comparisons test; n=9 *NPF*^*gut*^>, n=8 *NPF*^*gut*^>*TrpA1*, n=9 *NPF*^*gut*^>*TrpA1, NPFi*^*TRiP*^. (M) Survival under starvation of mated females. Kaplan-Meier log-rank tests. (N) TAG levels in mated females in the fed condition and following 24 hours’ starvation. Mann-Whitney U test; n=8 independent biological replicates with 3 animals per sample of fed *EEC*>, n=10 fed *EEC*>*NPFi*^*KK*^, n=9 starved *EEC*>, n=9 starved *EEC*>*NPFi*^*KK*^. Error bars indicate standard error of the mean (SEM). ns, non-significant; *, *p*<0.05; **, *p*<0.01; and ***, *p*<0.001.

To attribute these effects specifically to EEC-derived NPF, we measured the expression of *NPF* in dissected midguts and central nervous systems (CNS; brain and ventral nerve cord, VNC). *NPF* transcript levels were strongly reduced in the female midgut when *EEC*> was used to drive knockdown of *NPF*, whereas expression in the CNS was unaltered (Fig. 1E). We next examined the effectiveness and anatomical specificity of RNAi knockdown using immunostaining of female tissues. As expected, NPF immunofluorescence was eliminated in the EECs but not the CNS of *EEC*>*NPFi* animals (Fig. 1F and 1G). Furthermore, we confirmed that knockdown of *NPF* using a second RNAi line completely abolished midgut NPF immunofluorescence (Fig. S1F and S1G). Taken together this suggests that EEC-specific loss of *NPF* is responsible for the observed feeding phenotypes. To further support this we used *NPF::2A::GAL4* (*NPF*>), a CRISPR-mediated insertion of *T2A::GAL4* into the native *NPF* locus that drives GAL4 expression only in NPF-producing cells ^24^. Since NPF is expressed in both neurons and EECs, we used pan-neuronal *R57C10-GAL80*, an optimized *nSyb-GAL80* variant that suppresses neuronal GAL4 ^18^, to suppress the GAL4 activity of *NPF*> in the nervous system. We confirmed that this driver combination (*R57C10-GAL80; NPF*>), referred to hereafter as *NPF*^*gut*^>, efficiently knocks *NPF* transcript levels down in the midgut without affecting CNS expression (Fig. S1H), indicating that this driver specifically targets EEC-derived NPF. Consistent with this, NPF immunostaining in the CNS was unaffected when this driver was used to knock down *NPF*, while NPF signal in the EECs was abolished (Fig. 1H and S1I). Knockdown of *NPF* using this gut-NPF-specific driver caused a marked increase of sugar+yeast food intake over 30 minutes and sugar-only medium over 6 hours (Fig. 1I and S1J), similar to the phenotypes obtained with *EEC*>-driven loss of *NPF* in the EECs. Taken together these results indicate that gut endocrine cells produce NPF as a satiety signal that inhibits sugar consumption.

To examine the ability of circulating NPF to promote satiety, we injected synthetic NPF peptide into mated females. EEC-specific knockdown of *NPF* with two different RNAi lines induced hyperphagia, which in both cases was blocked by NPF injection (Fig. 1J and S1K). NPF injection did not affect hemolymph sugar levels (Fig. 1K), indicating that NPF can act systemically via the circulation to suppress food intake and, furthermore, that this effect is not a consequence of alterations in glycemic levels. Next, we expressed the thermosensitive Transient receptor potential A1 (TrpA1) cation channel ^25^ in the NPF^+^ EECs to enable induction of NPF release. Incubation at 29 °C, which induces TrpA1-mediated peptide release, inhibited food intake, an effect that was abolished by simultaneous *NPF* knockdown (Fig. 1L). Together these results suggest that NPF from these EECs is both necessary and sufficient to inhibit food intake and prevent food overconsumption.

Feeding behaviors are tightly coordinated with physiology in order to maintain metabolic balance. Our findings indicate that NPF acts as a satiety signal, which suggests that it should act after a meal. In this scenario NPF would be expected to promote storage and inhibit mobilization of energy. We therefore asked whether EEC-derived NPF affects energy metabolism. As an indirect measure of energy storage and mobilization, we assessed the starvation resistance of females with EEC-specific loss of *NPF*. Animals that have decreased energy stores or that mobilize their stores more quickly survive shorter during starvation. *NPF* knockdown in the EECs throughout development (using *EEC*> without *GAL80*^*TS*^) led to a decrease in starvation resistance in females but not males (Fig. 1M and S2A), in line with a recent study linking gut NPF to metabolic programs associated with energy storage ^19^. We confirmed this phenotype with adult-restricted knockdown (*EEC*> with *GAL80*^*TS*^), using two independent RNAi lines (Fig. S2B and S2C), suggesting that *NPF* knockdown affects energy metabolism. We next measured energy stores directly by assaying triacylglyceride (TAG) and glycogen content. Consistent with their shortened starvation survival, we found that females with EEC knockdown of *NPF* throughout development showed a marked decrease in both TAG and glycogen levels, while TAG levels were not affected by EEC-specific *NPF* loss in males (Fig. S2D). Adult-EEC-specific knockdown of *NPF* led to decreased TAG levels compared to the control only after 24-hour starvation (Fig. 1N and S2E). This suggests that while NPF does affect metabolism in the adult stage, it also has developmental effects that affect the adult. Together our findings indicate that a main function of EEC-derived NPF in the adult stage is the regulation of feeding, particularly the inhibition of sugar intake in females. We therefore reasoned that manipulations of gut NPF might affect circulating sugar levels, especially after a meal. To test this, we measured hemolymph glucose levels before and after food intake. Consistent with the increased post-starvation consumption exhibited by females with EEC suppression of *NPF*, we found that hemolymph glucose levels increased more after refeeding in these animals than in controls (Fig. S2F). Although animals lacking EEC-derived NPF were hypoglycemic after starvation, they reattained normal hemolymph sugar concentrations after 6 hours of re-feeding, consistent with their hyperphagia and showing that the mechanisms of sugar absorption and transport into circulation are functional. Dietary carbohydrates are important energy sources, but females require dietary protein for egg production. Our results indicate that mated females require gut NPF to adapt their feeding behavior and suppress sugar consumption.

### EEC-derived NPF suppresses sugar intake and promotes protein feeding

Our findings suggest that NPF acts as a sugar-satiety signal that balances carbohydrate intake with metabolic needs. To test this hypothesis directly, we examined whether gut NPF affects preference for dietary sugar when animals were given the choice between two different (1% and 10%) sucrose concentrations, based on feeding behaviors recorded with a FLIC assay. Females exhibited increased total feeding time on 10% sugar, and this preference was strongly enhanced by *NPF* knockdown in the EECs (Fig. 2A). We next measured sugar preference based on consumption in a two-choice CAFÉ assay, when flies were given the choice between ingesting 1% and 10% sucrose. Flies with EEC-specific *NPF* knockdown strongly preferred 10% sucrose compared to controls during both the initial 6 hours and over a longer time (Fig. 2B and S3A). We confirmed this effect using the *NPF*^*gut*^> driver together with an independent RNAi line (Fig. 2C and S3B). Together these results indicate that NPF is part of a post-ingestion sugar-sensing mechanism required to decrease sugar appetite.

**Figure 2.**
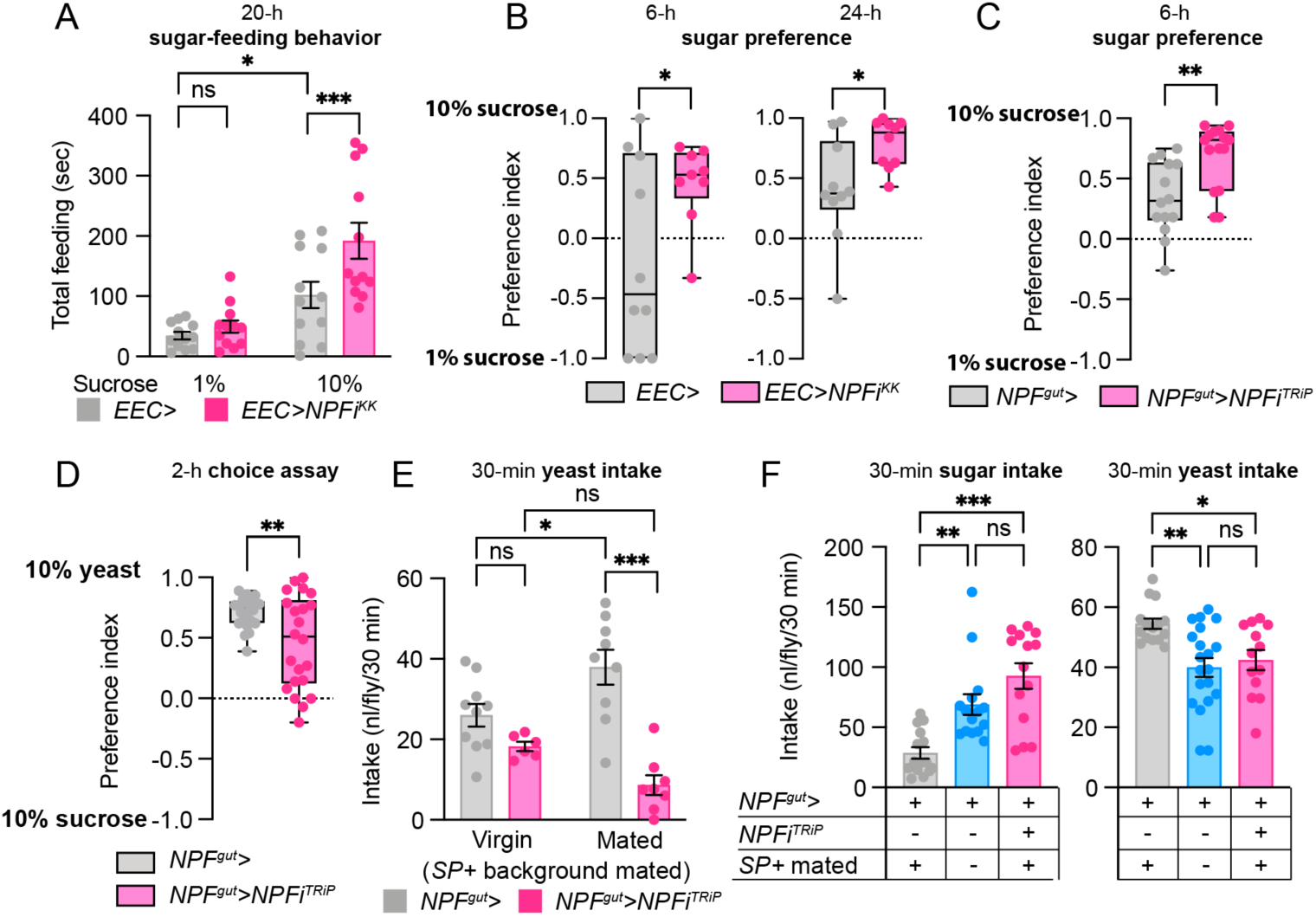
Gut-derived NPF regulates sugar intake and metabolism in females. (A) Time spent feeding on 1% or 10% sucrose over 6 hours of 15-hour-starved mated females measured using FLIC. One-way ANOVA with Bonferroni’s multiple comparisons test; all n=12. (B and C) Preference of 15-hour-starved female flies for 1% or 10% sugar measured over 6 or 24 hours by CAFÉ assay. (B) 6-hour preference: Student’s t-test; n=10 *EEC*>, n=9 *EEC*>*NPFi*^*KK*^. 24-hour preference: Student’s t-test; n=10 each. (C) Mann-Whitney U test; n=14 *NPF*^*gut*^>, n=15 *NPF*^*gut*^>*NPFi*^*TRiP*^. (D) Preference of 15-hour-starved mated female flies for 10% sucrose or 10% yeast over 2 hours measured by two-choice feeding assay. Student’s t-test; n=24 *NPF*^*gut*^>, n=22 *NPF*^*gut*^>*NPFi*^*TRiP*^. (E) Consumption of 10% yeast food over 30 minutes measured by dye assay in virgin and mated 15-hour-starved female flies. One-way ANOVA with Tukey’s post hoc test; n=10 *NPF*^*gut*^> virgins, n=6 *NPF*^*gut*^>*NPFi*^*TRiP*^ virgins, n=9 *NPF*^*gut*^> mated females, n=8 *NPF*^*gut*^>*NPFi*^*TRiP*^ mated females. (F) Intake of 10% sucrose (left panel) or 10% yeast (right panel) by 15-hour-starved females mated by SP-producing or SP-deficient males (*SP*^*0*^/*Df(3L)delta130*). Sucrose: Kruskal-Wallis with Dunn’s multiple comparisons test; n=15 *NPF*^*gut*^> females mated to *SP*^*+*^ males, n=14 *NPF*^*gut*^> females mated to *SP*-mutant males, n=15 *NPF*^*gut*^>*NPFi*^*TRiP*^ females mated to *SP*^*+*^ males. Yeast: one-way ANOVA with Dunnett’s multiple comparison test; n=15 *NPF*^*gut*^> females mated to *SP*^*+*^ males, n=20 *NPF*^*gut*^> females mated to *SP*-mutant males, n=13 *NPF*^*gut*^>*NPFi*^*TRiP*^ females mated to *SP*^*+*^ males. Error bars indicate standard error of the mean (SEM). ns, non-significant; *, *p*<0.05; **, *p*<0.01; and ***, *p*<0.001.

Given that females increase their preference for protein-rich food after mating to sustain metabolic requirements for egg production ^6^, we speculated that gut NPF might be important to reduce sugar appetite and increase preference for protein-rich food. To investigate whether NPF affects this preference, we used a two-choice food assay to measure preference between sugar-only or protein-rich yeast-only food after 3 days of protein deprivation, which increases preference for yeast food in mated females ^6^. Animals with EEC-specific *NPF* loss displayed a reduced preference for yeast food (Fig. 2D). This indicates that EEC-derived NPF, while inhibiting sugar intake, promotes yeast consumption, thereby switching food preference towards protein-rich food. In mated females, SP induces an increased preference for yeast, and this peptide has also been shown to potentiate NPF release from the midgut ^26^. We therefore hypothesized that NPF could be involved in mediating the SP-induced switch towards increased protein consumption in mated females. To test this possibility, we knocked *NPF* down in the EECs and examined whether it affected dietary yeast intake after mating in females, using the short-term dye-consumption assay with yeast-only food. As expected, control females increased their yeast consumption after mating with SP^+^ males (Fig. 2E). Virgin females lacking NPF consumed an amount of yeast similar to virgin control animals, and, consistent with our conjecture, mating by SP^+^ males did not induce these females to increase their yeast intake. To further demonstrate that loss of *NPF* in the EECs decreases consumption of dietary yeast in mated females, we measured long-term yeast consumption using the CAFÉ assay. As described above, mated females with EEC knockdown of *NPF* consume more sugar (Fig. 1D and 1I). Conversely when left with only the choice to consume yeast these animals consumed less food, confirming that loss of *NPF* reduces the intake of dietary yeast (Fig. S3C). Taken together our data show that mated females lacking EEC-derived NPF exhibit increased sugar consumption and decreased ingestion of yeast, suggesting that NPF from the midgut is necessary to induce preference towards yeast in females after mating. Upon mating, SP transferred from the male triggers this behavioral switch ^6^. Consistent with the notion that NPF is required for the mating-induced shift towards protein preference, females with EEC-specific *NPF* knockdown mated to SP^+^ males displayed a consumption pattern similar to that of control females mated to SP-mutant males – they consumed more sugar and less protein than females mated to SP^+^ males (Fig. 2F). Thus, females up-regulate their protein intake after mating in a SP-dependent manner, and our results indicate that NPF from the EECs is required for this shift in food choice.

### Sugar transporter 2 in NPF^+^ enteroendocrine cells governs food choice and release of nutrient-responsive NPF

Given these results that indicate that NPF acts as a mediator of sugar satiety, we asked whether NPF-producing EECs are activated by sucrose ingestion. In our initial analysis of a collection of RNAi lines (Fig. 1A), we identified *sugar transporter 2* (*sut2*), whose knockdown in the EECs of mated females increased their sugar feeding behavior (Fig. 3A). *Sut2* is expressed in adult EECs, according to the FlyGut-EEC single-cell transcriptomic resource ^27^, and it belongs to the glucose-transporter class of solute carrier (SLC) proteins, which in mammals are involved in the sugar-stimulated intestinal secretion of GLP-1 ^28^. SLC transporters are also involved in sugar-induced stimulation of hormonal release from EECs of *Drosophila*. The Sut2 paralog Sut1 has been linked to intestinal NPF secretion ^19^, and Glucose transporter 1 (Glut1) is involved in Burs secretion from EECs ^20^. Since we observed that EEC-specific knockdown of *sut2* phenocopied *NPF* knockdown by increasing sugar feeding, we hypothesized that NPF secretion from EECs might be regulated by Sut2. To validate this role of Sut2, we knocked down its expression specifically in NPF^+^ EECs and found that females displayed increased sugar feeding behavior similar to that observed in animals with knockdown of *NPF* itself (Fig. 3B, 3C, and S3D), suggesting that Sut2 might be required in NPF^+^ EECs as part of a mechanism governing NPF production or release.

**Figure 3.**
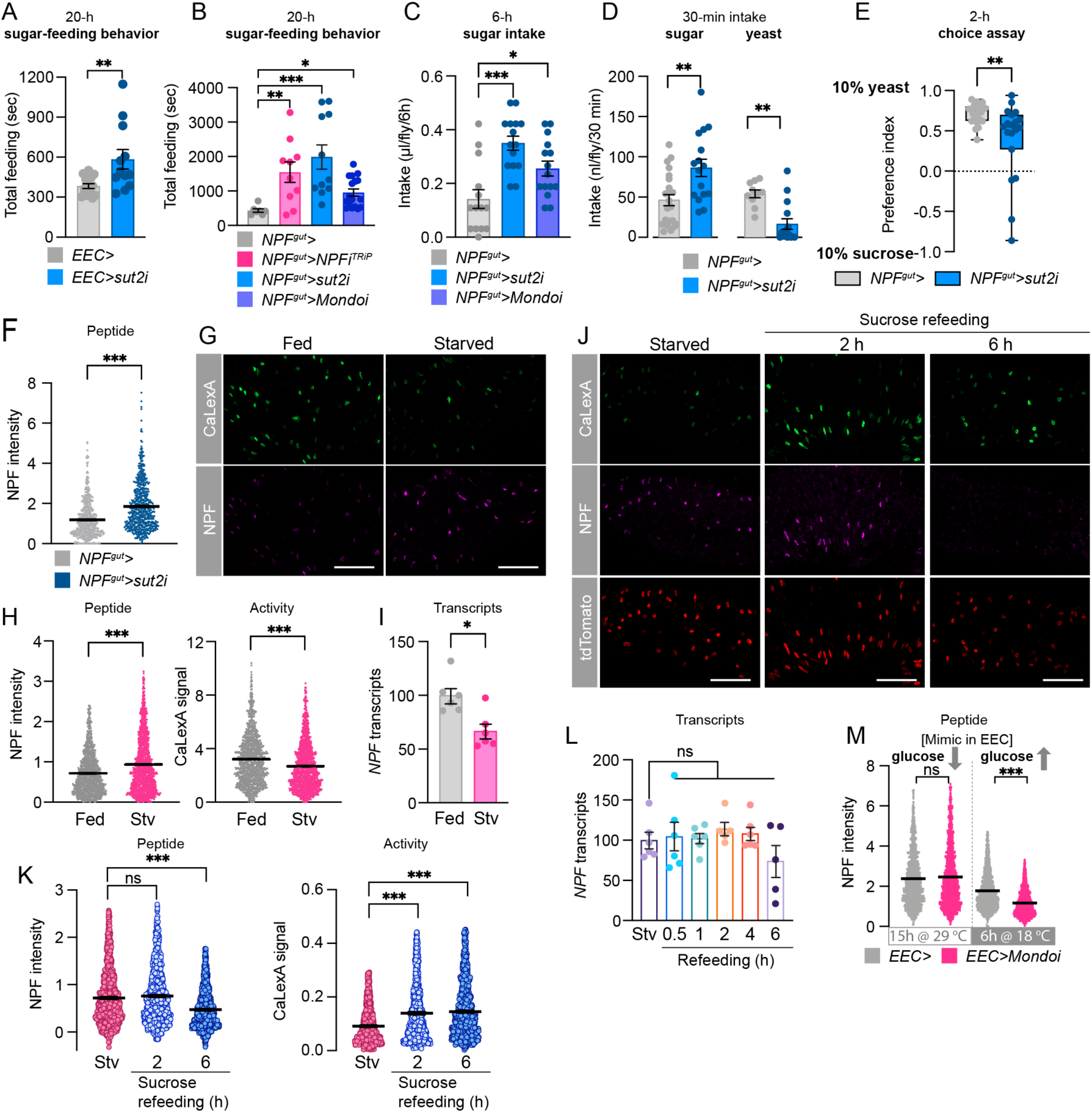
Sugar transporter 2 in the EECs regulates glucose-stimulated NPF secretion in mated females. (A and B) Time spent feeding on 10% sucrose over 24 hours in mated females measured using FLIC. (A) Student’s t-test; n=16 *EEC*>, n=9 *EEC*>*sut2i*. (B) One-way Kruskal-Wallis nonparametric ANOVA; n=7 *NPF*^*gut*^>, n=10 *NPF*^*gut*^>*NPFi*^*TRiP*^, n=11 *NPF*^*gut*^>*sut2i*, n=17 *NPF*^*gut*^>*Mondoi*. (C) Consumption of 10% sugar by 15-hour-starved mated females measured by CAFÉ assay over 6 hours. One-way ANOVA with Dunnett’s post-hoc test; n=14 *NPF*^*gut*^>, n=15 *NPF*^*gut*^>*sut2i*, n=15 *NPF*^*gut*^>*Mondoi*. (D) Consumption of 10% sucrose or 10% yeast by 15-hour-starved mated females, measured by dye assay over 30 minutes. Sucrose: Student’s t-test; n=22 *NPF*^*gut*^>, n=16 *NPF*^*gut*^>*sut2i*. Yeast: Mann-Whitney U test; n=10 *NPF*^*gut*^>, n=15 *NPF*^*gut*^>*sut2i*. (E) Preference of 15-hour-starved mated females for 10% yeast versus 10% sucrose, measured over 2 hours by two-choice consumption assay. Mann-Whitney U test; n=24 *NPF*^*gut*^>, n=19 *NPF*^*gut*^>*sut2i*. (F) Midgut NPF staining intensity in mated females with *sut2* knockdown in NPF^+^ EECs. Mann-Whitney U test; n=571 cells from 6 guts for *NPF*^*gut*^>, n=625 cells from 5 guts for *NPF*^*gut*^>*sut2i*. (G) Representative images of midgut NPF-cell activity measured by calcium (CaLexA) reporter system (*NPF*>*LexA::NFAT::VP16; LexAop-GFP*) and NPF staining intensity in fed and 24-hour-starved mated females. Scale bars, 50 µm. (H) Quantification of G. Mann-Whitney U tests; for NPF intensity, n=1167 cells from 8 fed guts, n=1394 cells from starved 8 guts; for CaLexA intensity, n=1209 cells from 9 fed guts, n=1405 from 9 starved guts. (I) *NPF* transcripts in midguts of fed and 24-hour-starved mated females. Mann-Whitney U test; n=5-6 samples of 5 guts each, per condition. (J) Representative images of NPF staining intensity and NPF-cell activity in 24-hour-starved, 2-hour-sugar-refed, and 6-hour-sugar-refed mated females measured by modified CaLexA calcium reporter system in which all cells within the GAL4 pattern, active or not, are marked with tdTomato (*NPF-GAL4; UAS-tdTomato; UAS-LexA::NFAT::VP16; LexAop-GFP*). Scale bars, 50 µm. (K) Quantification of J. Kruskal-Wallis nonparametric ANOVA. Eight guts per condition; NPF intensity: n=1297 starved cells, n=1038 2-h-refed cells, n=1216 6-h-refed cells. CaLexA intensity: n=1169 starved cells, n=954 2-h-refed cells, n=1151 6-h-refed cells. (L) *NPF* transcript levels in the midguts of mated females refed for varying periods after 24-hour starvation (stv). One-way ANOVA with Dunnett’s post-hoc test; n=6 replicates of 6 midguts for stv, refed 0.5 h, 1 h, and 4 h; n=5 refed 2 h and 6 h. (M) NPF staining intensity in midguts from mated females with 15 hours’ activation of EEC knockdown of *Mondo* (transfer to 29 °C to inactivate GAL80^TS^) and following 6-hour re-activation of Mondo expression (transfer to 18 °C to renature GAL80^TS^). Mann-Whitney U test; 15 h at 29 °C: n=12 *EEC*>, n=14 *EEC*>*Mondoi;* 6 h at 18 °C: n=17 *EEC*>, n=17 *EEC*>*Mondoi*. Error bars indicate standard error of the mean (SEM). ns, non-significant; *, *p*<0.05; **, *p*<0.01; and ***, *p*<0.001.

In *Drosophila* and mammals, the sugar-sensitive transcription factor Mondo/ChREBP (Carbohydrate-responsive-element-binding protein) contributes to many of the cellular responses to sugar ^29^. To probe the molecular sugar-sensing mechanisms regulating NPF, we silenced *Mondo* specifically in NPF^+^ EECs and found that this manipulation increased both sugar feeding behavior and sugar intake, although not as dramatically as loss of *NPF* or *sut2* (Fig. 3B and 3C). This suggests that although other mechanisms are likely involved, Mondo/ChREBP-mediated sugar sensing may contribute to NPF regulation in EECs.

Because *NPF* loss increased sugar intake and decreased protein feeding, we investigated whether Sut2 also affects sugar versus protein intake. In this experiment, we measured mated females’ consumption of either 10% sucrose or 10% yeast medium when presented separately, and we also assessed their dietary preference in an assay in which they were given a choice between these two foods. We found that knockdown of *sut2* in NPF^+^ EECs led to a strongly increased intake of sucrose and a strong decrease in consumption of yeast (Fig. 3D). When given a choice between these two foods, animals lacking *sut2* in NPF^+^ EECs exhibited a reduced preference for dietary yeast (Fig. 3E). These results show that loss of *sut2* in NPF^+^ EECs shifts dietary preference from protein-rich food towards sugar, similar to knockdown of *NPF* in these gut cells. To examine whether Sut2 regulates NPF production and secretion, we measured midgut *NPF* expression and protein levels in animals with knockdown of *sut2* in NPF^+^ EECs. We found that this cell-specific knockdown of *sut2* led to strong accumulation of NPF peptide (Fig. 3F), even though *NPF* transcript levels were reduced (Fig. S3E), suggesting that *sut2* loss in NPF^+^ EECs leads to retention of NPF and thus that Sut2 is required for normal NPF release. We also found that *sut2* transcript levels in the entire midgut were strongly reduced (∼80%) by knockdown targeted only to the NPF^+^ endocrine gut cells (Fig. S3E), demonstrating that *sut2* is predominantly expressed in NPF^+^ EECs.

To assess more directly whether sugar regulates EEC activity levels and thereby governs *NPF* expression or secretion, we compared animals fed sugar+yeast food, animals starved for 24 hours, and animals refed with sucrose only after 24-hour food deprivation. We observed cellular calcium-signaling history using the CaLexA reporter system ^30^, and we measured midgut *NPF* transcript and intracellular peptide levels. After 24 hours of starvation, we observed decreased calcium signaling in the NPF^+^ EECs (Fig. 3G and 3H), indicating a reduction in their activity. NPF protein levels were increased in the EECs of animals starved 24 hours, although *NPF* transcript levels were reduced (Fig. 3G-3I). Together, these data suggest a reduction in NPF^+^ EEC activity and thus lower peptide secretion under starvation. Refeeding with sucrose after starvation elicited a strong increase in calcium signaling within two hours in the NPF^+^ EECs, associated with a decrease in NPF peptide staining within six hours of refeeding (Fig. 3J, 3K, and S3F). Since midgut *NPF* transcript levels were unaltered under these conditions (Fig. 3L), these results indicate that midgut NPF^+^ cells are activated by dietary sugar, leading to their secretion of NPF peptide.

We then used genetic methods to mimic sugar sensing occurring in the EECs following a meal. We first induced RNAi-mediated silencing of *Mondo/ChREBP* by switching flies to 29 °C to inactivate GAL80^TS^ for 15 hours, to reduce sugar sensing in the EECs. We then reactivated Mondo/ChREBP signaling, to mimic sugar-sensing occurring after a meal, by switching animals from 29 °C back to 18 °C to renature GAL80^TS^ and thereby inactivate the RNAi effect. Reactivation of Mondo/ChREBP signaling in the EECs caused a decrease in NPF peptide levels in these cells (Fig. 3M) without altering *NPF* expression (Fig. S3G), indicating increased NPF secretion, consistent with the notion that sugar sensing in the EECs is associated with NPF release. Taken together our findings indicates that sugar intake leads to increased calcium levels in NPF^+^ EECs, which leads to NPF release. Furthermore, this release of NPF requires the glucose-transporter-family protein Sut2, and this process, at least in part, involves Mondo/ChREBP-mediated sugar sensing.

### NPF suppresses energy mobilization in adipose tissues via NPFR

We then asked which target tissues might be involved in NPF-mediated appetite regulation. To determine whether the effects of gut NPF are mediated by receptive cells of the nervous system, we knocked *NPFR* down either pan-neuronally or globally in the entire organism, using *elav-GAL4* or *daughterless (da)-GAL4*. Notably, whereas EEC-specific *NPF* knockdown induced overfeeding (Fig. 1D), adult females with neuronal knockdown of *NPFR* showed decreased food intake (Fig. 4A), consistent with previous reports that the neuronal NPF/NPFR signaling promotes feeding ^31^ and indicating that other tissues mediate the downregulation of sugar intake induced by gut NPF. Consistent with this, we found that global knockdown of *NPFR* led to a feeding phenotype intermediate between those observed with gut or neuronal *NPF* signaling loss (Fig. 4A), likely reflecting opposing effects on feeding of NPF signaling within different organs.

**Figure 4.**
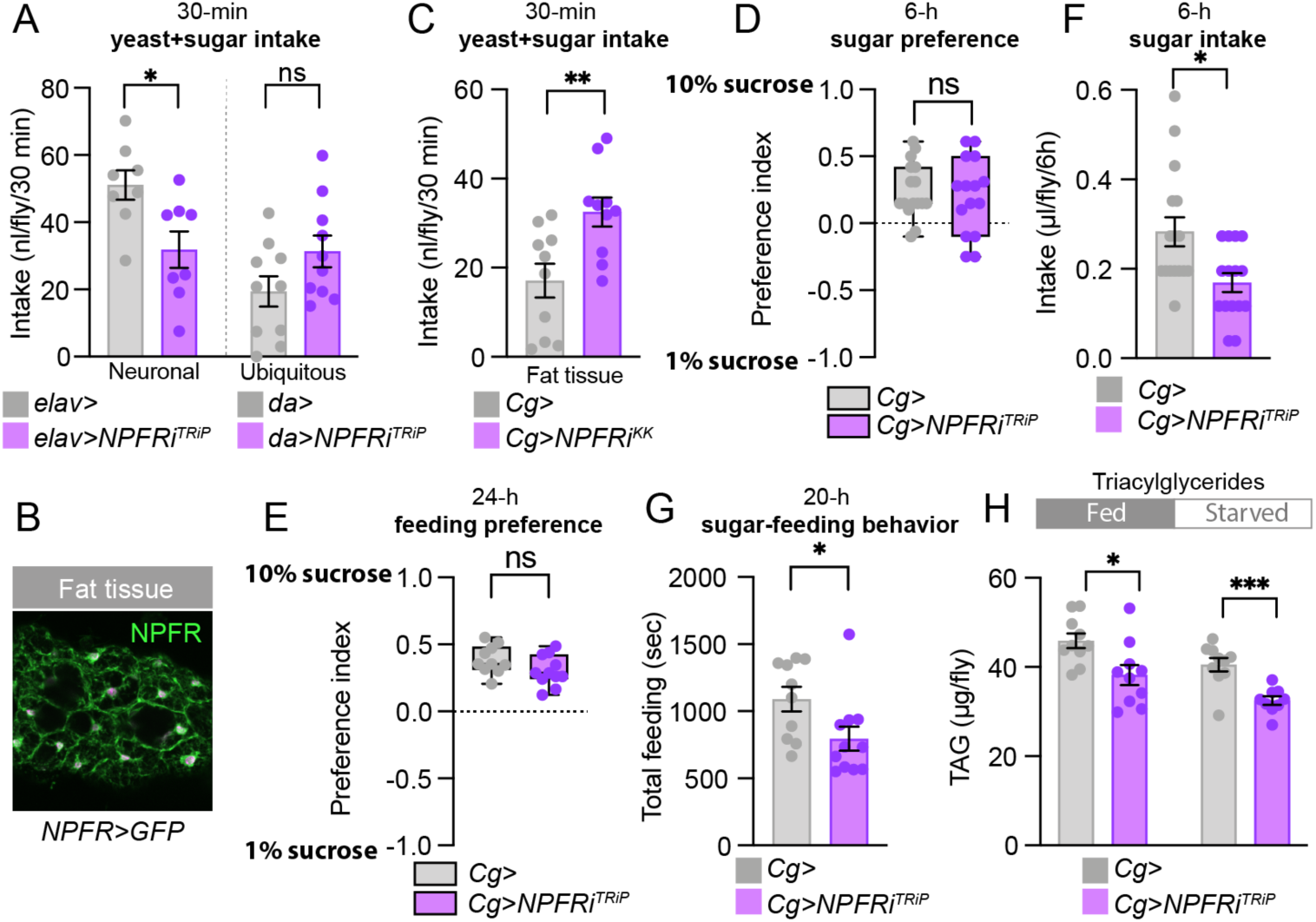
Loss of NPFR in the fat affects metabolism but does not increase preference for dietary sugar in mated females. (A) Food intake of 15-hour-starved mated females over 30 minutes in animals with knockdown of *NPFR* in the nervous system (*elav*>) or the entire body (*da*>) measured by dye assay with sugar+yeast food (9% sugar and 8% yeast). Student’s t-test; n=10, except n=8 for *elav*>. (B) Fat-body immunostaining of *NPFR*>*mCD8::GFP* reporter in mated females. (C) Food intake of 15-hour-starved mated females measured by dye assay with sugar+yeast food (9% sugar and 8% yeast) in fat-body *NPFR*-knockdown animals. Student’s t-test; n=10 *Cg*> or *Cg*>*NPFRi*^*KK*^. (D) Consumption preference of 15-hour-starved mated females for 1% or 10% sucrose, measured over 6 hours by CAFÉ assay. Student’s t-test, n=16 *Cg*>, n=15 *Cg*>*NPFRi*^*TRiP*^. (E) Preference of mated females for 1% *versus* 10% dietary sucrose measured by FLIC over 24 hours. Student’s t-test; n=10 *Cg*>, n=11 *Cg*>*NPFRi*^*TRiP*^. (F) Sucrose intake of 15-hour-starved mated females measured over 6 hours by CAFÉ assay. Mann-Whitney U test; n=15 for *Cg*>, n=16 *Cg*>*NPFRi*^*TRiP*^. (G) Time spent feeding on sucrose of mated females measured over 24 hours using FLIC. Mann-Whitney U test (n=10 *Cg*>, n=11 *Cg*>*NPFRi*^*TRiP*^). (H) Triacylglyceride (TAG) levels in fed and 15-hour-starved fat-body *NPFR*-knockdown animals. Fed: Student’s t-test; n=10 *Cg*> or *Cg*>*NPFRi*^*TRiP*^. Starved: Mann-Whitney U test; n=10 *Cg*>, n=9 *Cg*>*NPFRi*^*TRiP*^. Error bars indicate standard error of the mean (SEM). ns, non-significant; *, *p*<0.05; **, *p*<0.01; and ***, *p*<0.001.

To assess receptor expression in other target tissues, we used a CRISPR-mediated knock-in of *T2A::GAL4* into the native *NPFR* locus (*NPFR::T2A::GAL4*, hereafter *NPFR*>) ^32^ to express *UAS-mCD8::GFP* for visualizing receptor expression. We observed GFP expression in the fat body (Fig. 4B), a tissue analogous to the adipose and liver in mammals, and we therefore next suppressed *NPFR* expression in this tissue using *Cg-GAL4* (*Cg*>) to investigate whether fat-body NPFR signaling regulates sugar appetite. Although fat-body-specific *NPFR* knockdown in adult females did lead to increased short-term (30-minute) intake of food containing both sugar+yeast (Fig. 4C and S4A), animals lacking *NPFR* expression in the fat tissue did not display increased preference for feeding on high sugar concentrations over 6 hours or 24 hours (Fig. 4D and 4E). Indeed, suppression of *NPFR* in the fat body caused a decrease in sucrose intake over 6 hours and reduced sugar feeding behavior over 24 hours (Fig. 4F and 4G). Thus, fat-body NPFR signaling does not appear to underlie the specific feeding phenotypes observed with gut *NPF* loss. We next asked whether fat-body NPFR mediates the effect of gut NPF on metabolism. Like animals with gut-specific *NPF* knockdown, fat-body *NPFR* knockdown animals were more sensitive to starvation (Fig. S4B). Furthermore, fat-body knockdown of *NPFR* led to a metabolic phenotype similar to that seen with loss of gut NPF, including reduced whole-body TAG and glycogen levels (Fig. 4H and S4C). Thus, loss of *NPFR* function in the fat body recapitulates the metabolic effects, but not the increased sugar preference, observed in mated female flies with EEC-specific *NPF* knockdown. These findings indicate that gut-derived NPF acts on NPFR in the fat body as part of a metabolic pathway that maintains energy homeostasis.

### NPF suppresses glucagon-like signaling to restrict sugar intake and drive hunger for protein food

Our experiments imply that gut NPF signaling regulates sugar appetite via tissues other than the CNS and fat body. In *Drosophila*, the brain cells that produce insulin express NPFR ^19^, and these cells also regulate aspects of feeding and satiety ^33^. However, these cells should be targeted by pan-neuronal knockdown of *NPFR* (Fig. 4A), which did not recapitulate the feeding phenotype of gut *NPF* loss, suggesting that gut-derived NPF does not act through insulin to modulate preference for dietary sugar and protein. To identify the tissue mediating this effect, we examined *NPFR* expression in other tissues, which revealed expression of the receptor in the AKH-producing cells (APCs), part of a neuroendocrine tissue called the corpora cardiaca (CC) (Fig. 5A). During fasting or exercise, which tend to reduce blood-sugar levels, glucagon signaling acts via the adipose and hepatic tissues to promote energy mobilization and thus to prevent hypoglycemia ^2^. In flies, glucagon-like AKH is released during starvation and acts through its receptor, AkhR, in the fat body to promote the mobilization of stored energy and is also thought to act as a hunger signal to drive feeding behaviors ^18,34-37^. However, whether AKH regulates sugar- or protein-specific feeding is unknown. We hypothesized that gut-derived NPF, released in response to sugar feeding, might suppress AKH release from the APCs in the fed state. Consistent with a recent report ^19^, we found that knocking down *NPFR* in the APCs using *AKH-GAL4* (*AKH*>) resulted in decreased AKH peptide levels within these cells in fed mated females (Fig. 5B), indicating that NPFR is required to suppress AKH release when the animal has ingested food. Indeed, AKH levels in the APCs of fed flies with *NPFR* knockdown were reduced to levels similar to those observed in control animals after starvation (Fig. 5B), which induces AKH release. In agreement with the notion that NPF acts specifically upon feeding to suppress AKH release, and not in starved conditions, *NPFR* loss in the APCs did not affect levels of AKH in starved animals. AKH promotes lipid and glycogen breakdown in the fat body, and we therefore tested whether NPFR activity in the APCs regulates metabolism. As with knockdown of *NPF* in the midgut, adult females with knockdown of *NPFR* in the APCs showed reduced TAG and glycogen levels and increased susceptibility to starvation, consistent with an increase in AKH signaling (Fig. S5A and S5B).

**Figure 5.**
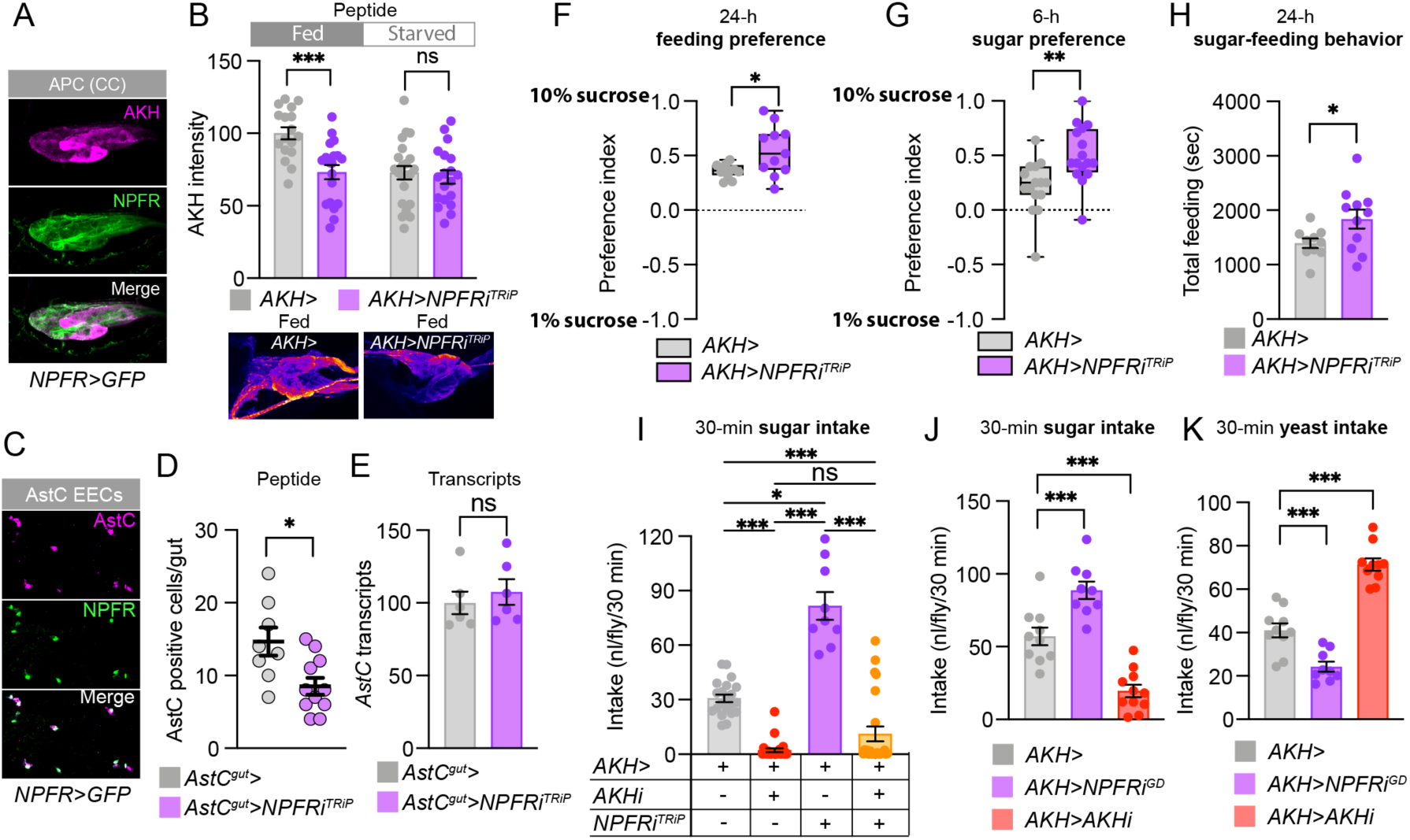
Knockdown of *NPFR* in the AKH-producing cells of the CC phenocopies gut *NPF*-loss feeding phenotypes in mated females. (A) Immunohistochemistry of AKH-producing cells (APCs) shows *NPFR*>*GFP* reporter expression in APCs of mated females. (B) Quantification of AKH levels in fed and 15-hour-starved mated females with *NPFR* knockdown in the APCs. Student’s t-test; n=17 fed *AKH*>, n=19 fed *AKH*>*NPFRi*^*TRiP*^, n=22 starved *AKH*>, n=19 starved *AKH*>*NPFRi*^*TRiP*^. (C) Immunohistochemistry of guts from *NPFR*>*GFP* mated female fllies showing co-localization of *NPFR* reporter and AstC. (D) Quantification of the number of cells per gut showing detectable AstC staining with and without *NPFR* knockdown in AstC^+^ EECs of fed mated females. Student’s t-test; n=8 *AstC*^*gut*^>, n=11 *AstC*^*gut*^>*NPFRi*^*TRiP*^. (E) *AstC* transcript levels quantified by qPCR from five midguts per each replicate from fed mated females. Student’s t-test; n=6 *AstC*^*gut*^>, n=6 *AstC*^*gut*^>*NPFRi*^*TRiP*^. (F) Preference of mated females for 1% and 10% sugar measured by FLIC over 24 hours. Student’s t-test; n=11 *AKH*>, n=11 *AKH*>*NPFRi*^*TRiP*^. (G) Consumption preference of 15-hour-starved mated females for 1% and 10% sucrose measured by CAFÉ assay over 6 hours. Student’s t-test; n=15 *AKH*>, n=16 *AKH*>*NPFRi*^*TRiP*^. (H) Time spent feeding on 10% sucrose of mated females measured by FLIC over 24 hours. Student’s t-test; n=10 *AKH*>, n=11 *AKH*>*NPFRi*^*TRiP*^. (I) Rescue of sugar overconsumption of 15-hour-starved mated females with *NPFR* knockdown in the APCs by simultaneous *AKH* knockdown measured over 30 minutes by dye assay with 10% sucrose medium. Kruskal-Wallis with Dunn’s test; n=21 *AKH*>, n=24 *AKH*>*AKHi*^*KK*^, n=9 *AKH*>*NPFRi*^*TRiP*^, n=24 *AKH*>*AKHi*^*KK*^, *NPFRi*^*TRiP*^. (J and K) Sucrose or yeast intake of 15-hour-starved mated females measured over 30 minutes by dye assay with 10% sucrose or 10% yeast medium. (J) One-way ANOVA; n=10 *AKH*>, n=9 *AKH*>*NPFRi*^*GD*^, n=11 *AKH*>*AKHi*^*KK*^. (K) One-way ANOVA; n=10 *AKH*>, n=9 *AKH*>*NPFRi*^*GD*^, n=10 *AKH*>*AKHi*^*KK*^. Error bars indicate standard error of the mean (SEM). ns, non-significant; *, *p*<0.05; **, *p*<0.01; and ***, *p*<0.001.

We recently reported that the peptide hormone AstC released by midgut EECs promotes AKH release during starved conditions ^18^. Since our results indicate that EEC-derived NPF acts to inhibit AKH secretion, we wondered whether NPF signaling might also inhibit AstC release from the midgut to suppress the activation of the AKH axis at multiple hierarchical levels. We first examined expression of NPFR reporter in the midgut, which revealed that NPFR is indeed expressed in AstC^+^ EECs (Fig. 5C), consistent with single-cell RNAseq data ^38^. To test whether NPF signaling directly inhibits release of AstC from these cells, we silenced *NPFR* expression specifically in AstC^+^ EECs using *AstC-GAL4* (*AstC*>) with pan-neuronal GAL80 (*R57C10-GAL80, AstC*>; *AstC*^*gut*^> hereafter) to suppress nervous-system GAL4 activity. This manipulation did not alter *AstC* expression, but it did lead to a reduction in the number of cells containing detectable AstC peptide (Fig. 5D and 5E), suggesting that loss of *NPFR* cell-autonomously increases AstC release. This increased EEC-derived AstC release, disinhibited through loss of *NPFR*, would be expected to promote AKH release, leading to more-rapid depletion of energy stores and therefore reduced capacity to survive starvation ^18^. To test this, we assessed starvation resistance in adult females with *NPFR* knockdown targeted to the AstC^+^ EECs, and indeed this knockdown led to a clear reduction in the capacity of these animals to survive starvation (Fig. S5C). Taken together our results indicate that in response to sugar intake, gut-derived NPF inhibits the AKH axis at three levels: blocking the release of adipokineticotropic AstC from midgut EECs, blocking the release of AKH itself from the APCs, and counteracting AKH’s effects on the fat body.

We then tested whether NPFR regulates sugar versus protein-specific feeding through NPFR in the APCs. Knockdown of *NPFR* in mated female APCs increased food intake and elevated sugar feeding and preference, determined by both sugar feeding behaviors and sugar consumption (Fig. 5F-5H and S5D-S5I), similar to animals with loss of *NPF* in the midgut. To determine whether AKH mediates the effects of *NPFR* loss on feeding, we examined the ability of *AKH* knockdown to rescue the sugar-overeating phenotype caused by *NPFR* RNAi in the APCs. We found that in mated females, knockdown of *AKH* completely abolished the sugar-overeating phenotype induced by loss of NPFR in the APCs (Fig. 5I), suggesting that AKH is the primary factor mediating the feeding effects of NPFR signaling.

Although previous reports have found that loss of AKH reduces food intake ^18,39^, the role of AKH in governing nutrient-specific feeding is completely unknown. To test the involvement of NPFR and AKH in balancing sugar versus protein feeding, we first assessed sugar consumption in adult females with APC-specific *NPFR* knockdown, using a second *NPFR-RNAi* construct. We again observed that *NPFR* knockdown led to increased sugar feeding, and we confirmed that *AKH* loss caused the opposite effect by reducing sugar intake (Fig. 5J). We then measured these animals’ consumption of protein-rich yeast food and found that *NPFR* knockdown in the APCs induced a clear reduction in yeast intake (Fig. 5K), similar to what we observed in animals with midgut-specific *NPF* loss (Fig. 2E and 2F). Furthermore, animals lacking *AKH* showed a marked increase in yeast consumption (Fig. 5K), indicating increased preference for protein-rich food. Together our results indicate that, in mated females, gut-derived NPF acts on the APCs via NPFR to inhibit AKH release after a sugar-rich meal, to suppress further sugar feeding while promoting protein intake.

### AKH regulates specific appetite for sugar and yeast food

AKH is described as a generic hunger hormone released during nutritional deprivation. However, our findings indicate that AKH has nutrient-specific effects, promoting sugar intake at the expense of protein consumption. To confirm this effect, we examined the feeding behavior of mated *AKH*-mutant females. These animals exhibited significantly reduced sugar intake (Fig. 6A), suggesting that AKH promotes sugar preference. To test this directly, we examined the food preferences of mated *AKH* mutant females. We found that loss of *AKH* either through knockdown or mutation led to a striking shift in preference towards yeast (Fig. 6B). This suggests that AKH is not simply a hunger hormone as previously reported ^18,34,40^, but rather has a more complex role in regulating nutrient-specific appetite for sugar versus protein-rich food.

**Figure 6.**
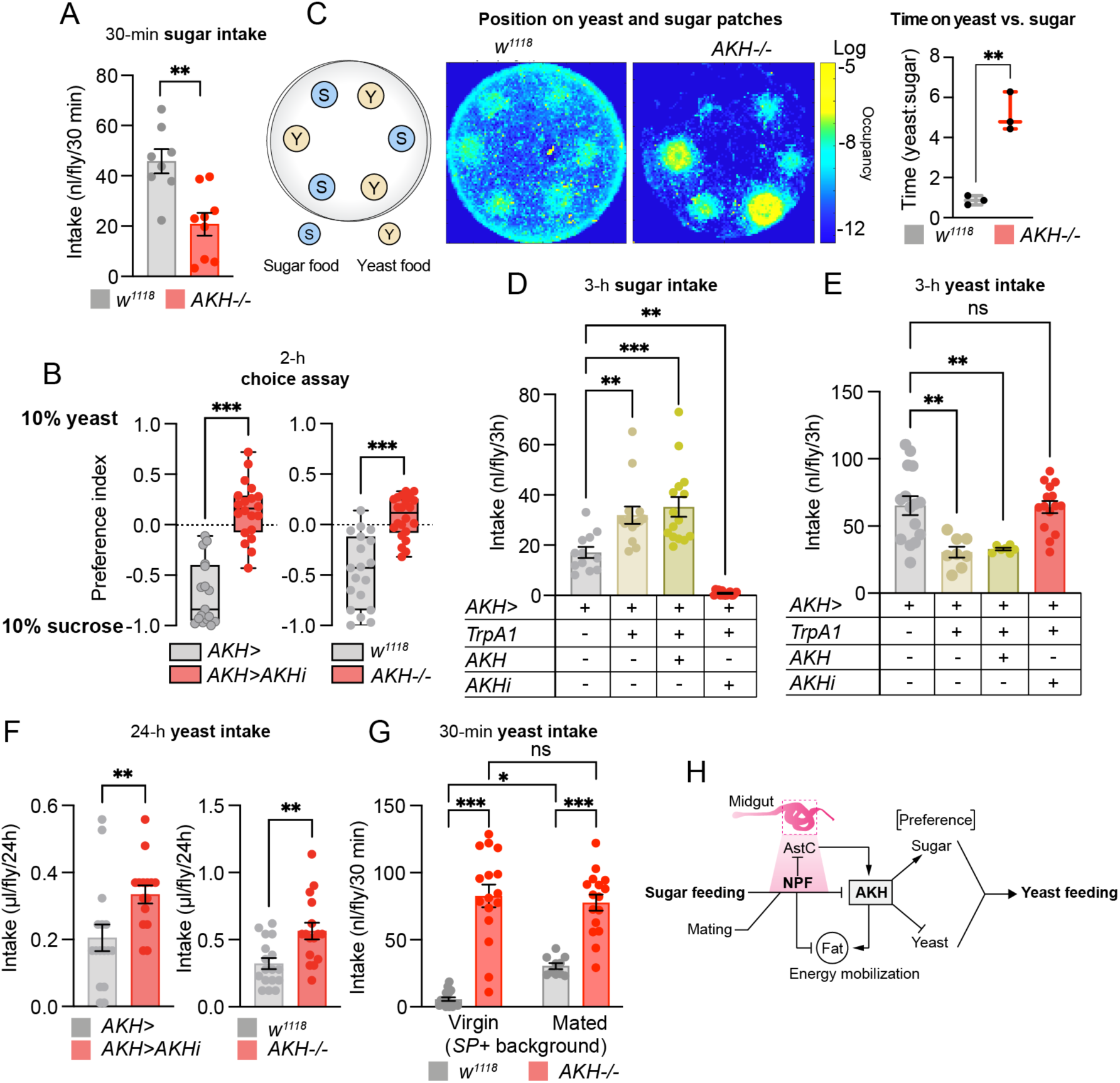
AKH promotes sugar feeding and suppresses protein intake in females. (A) Sucrose intake of 15-hour-starved mated females measured over 30 minutes by dye assay with 10% sucrose medium. Student’s t-test; sucrose food: n=8 *w*^*1118*^, n=9 *AKH*^-/-^. Yeast food: n=10 *w*^*1118*^, n=16 *AKH*^-/-^. (B) Consumption preference of 15-hour-starved mated females for 10% sucrose or 10% yeast measured over 2 hours using the two-choice assay. Mann-Whitney U test; n=17 *AKH*>, n=20 *AKH*>*AKHi*^*KK*^. Student’s t-test; n=19 *w*^*1118*^, n=22 *AKH*^-/-^. (C) Heat map of 30 minutes’ tracking of twelve 15-hour-starved mated females per genotype on sugar or yeast patches. Ratio of time spent on yeast *versus* sugar patches. Student’s t-test; n=3. (D and E) Sugar or yeast consumption measured over 3 hours in mated females using the dye assay, at 29 °C for TrpA1 activation. (D) One-way ANOVA; n=12 *AKH*>, n=14 *AKH*>*TrpA1*, n=15 *AKH*>*TrpA1+AKH*, n=15 *AKH*>*TrpA1+AKHi*). (E) One-way ANOVA; n=15 *AKH*>, n=8 *AKH*>*TrpA1*, n=7 *AKH*>*TrpA1+AKH*, n=15 *AKH*>*TrpA1+AKHi*. (F) Yeast intake of 15-hour-starved mated females measured over 24 hours by CAFÉ assay with 10% yeast extract medium. Left panel, *AKH* knockdown; right panel, *AKH* mutant. Student’s t-test; n = 16 *AKH*>, n = 15 *AKH*>*AKHi*^*KK*^, n = 17 *w*^*1118*^, n = 16 *AKH*^-/-^. (G) Consumption of 10% yeast food over 30 minutes measured by dye assay in virgin and mated 15-hour-starved female flies. One-way ANOVA with Tukey’s post hoc test; n=10 *w*^*1118*^ virgins, n=16 *AKH*^-/-^ virgins, n=10 mated *w*^*1118*^ females, n=16 mated *AKH*^-/-^ females. (H) A model of the NPF axis. Sugar feeding, potentiated by mating, promotes EEC NPF release. This NPF acts (1) on AstC^+^ EECs to block AstC release, thus indirectly blocking AKH release; (2) on the APCs, directly blocking AKH release; and (3) on the fat to block mobilization, counteracting the effects of AKH. AKH acts to drive feeding preferences towards sugar and away from protein. Error bars indicate standard error of the mean (SEM). ns, non-significant; *, *p*<0.05; **, *p*<0.01; and ***, *p*<0.001.

To further investigate the role of AKH in governing food-seeking behavior and preference towards specific nutrients, we used automated video tracking of individual flies in a foraging arena containing sugar-only and yeast-only food patches (Fig. 6C). Consistent with their increased intake and preference for yeast food, mated *AKH* mutant females displayed a strong increase in the amount of time spent exploring the yeast food, while control flies explored both types of food patches equally (Fig. 6C). This further supports a role for AKH in controlling feeding decisions, biasing behavior towards sugar intake while suppressing protein-specific hunger. To examine this role further, we induced peptide release from the APCs of fed animals using TrpA1. Activation of the APCs induced an increase in sugar intake (Fig. 6D), despite these animals’ fed state, along with a decrease in yeast intake (Fig. 6E), indicating that hormonal release from the APCs is sufficient to induce a feeding bias towards sugar and away from protein. Overexpression of AKH did not further enhance these phenotypes, suggesting a saturation of AKH signaling, while both effects were completely blocked by simultaneous *AKH* knockdown (Fig. 6D and 6E), indicating that the effect is mediated by AKH. Together these findings show that AKH is a hormone that controls selective feeding decisions by increasing appetite for sugar and reducing huger for protein food.

AKH has recently emerged as a key factor in sex-specific metabolic regulation ^41^. To determine whether AKH influences feeding in a sex-specific way, we analyzed yeast intake in control males and mated males lacking AKH. Unlike mated females, in which *AKH* loss promoted yeast intake (Fig. 6F), males lacking AKH exhibited a decrease in yeast intake, indicating that AKH plays a sexually dimorphic role in feeding decisions (Fig. S6). Together these findings indicate that, in mated females, gut-derived NPF inhibits AKH secretion, which suppresses sugar appetite and increases hunger for protein food. In this scenario, the increased AKH signaling in virgins is what promotes preference for dietary sugar over yeast food in the virgin state. We therefore conjectured that loss of AKH would increase yeast intake in virgin females to a level similar to that exhibited by mated females. As expected, we found that while control females increased their yeast consumption in response to mating, *AKH*-mutant virgins displayed a striking overconsumption of yeast food that was not altered by mating (Fig. 6G). We propose that sugar-induced AKH-suppressive NPF signaling from the gut constitutes a hormonal axis that switches feeding preference toward protein-rich food after mating in females (Fig. 6H).

## Discussion

To maintain nutritional homeostasis, animals need to match their ingestion of specific nutrients to their needs. This is achieved by modulating appetite towards the specific nutrients needed. A number of factors, including gut hormones, that regulate food consumption have been identified in both flies and mammals, and studies have also reported central brain mechanisms that induce ingestion of protein food in response to amino acids deprivation, sense amino acids and promote food consumption, and reject food lacking essential amino acids ^1,6,13,42,43^. However, little is known about the hormonal mechanisms that regulate nutrient-specific appetite, and gut hormones that regulate selective food intake are completely unknown. Our findings indicate that gut-derived NPF is a selective driver of sugar satiety and protein hunger, providing a basis for understanding these mechanisms. Gut-hormone-based therapies that inhibit appetite offer promising new directions for weight-loss treatments ^15^. For example, Fibroblast growth factor 21 (FGF21) is a liver-derived hormone that promotes protein consumption, and it is emerging as a promising target for metabolic disorders ^44^. Uncovering appetite-regulatory gut hormones like NPF that specifically inhibit sugar consumption while promoting intake of protein-rich food could provide effective new weight-management strategies, promoting healthier food choices and limiting the overconsumption of sugar, which can lead to obesity and metabolic disorders.

Our results suggest that Sut2, which belongs to the SLC2 family of sugar transporters, is required for NPF secretion from the EECs. Sut2 is the closest *Drosophila* homolog of human SLC2A7 (GLUT7), a transporter expressed mainly in the intestine whose function is poorly defined ^45^. In flies, GLUT1 is important for Burs secretion from the EECs, and Sut1, another SLC2-family sugar-transporter protein, was recently shown to be involved in midgut NPF release ^19,20^. Our results indicate that Sut2 is involved in controlling the release of NPF from EECs and is thus responsible, at least in part, for the mechanism by which NPF-mediated gut signaling controls feeding decisions. Indeed, we show that loss of *sut2* in NPF^+^ EECs leads to increased sugar consumption and decreased preference for protein-rich yeast food, phenocopying the loss of *NPF* itself in the gut and suggesting that Sut2 plays a central role in NPF^+^ EECs for nutrient-specific appetite regulation. In mammals, several mechanisms regulate glucose-stimulated GLP-1 secretion from intestinal endocrine cells, involving sodium-glucose cotransporter 1 (SGLT1), the glucose transporter GLUT2, and sweet taste receptors ^28^. Future studies should investigate whether GLUT7, like its *Drosophila* homolog Sut2, affects appetite-regulatory mechanisms in the mammalian gut.

NPF is orthologous with the mammalian NPY family of gut-brain peptides, including peptide YY (PYY), pancreatic polypeptide (PP), and NPY itself, that regulate food-seeking behaviors and metabolism ^46,47^. Like mammalian NPY-family hormones, *Drosophila* NPF is expressed in both the nervous system and the gut. While NPY is abundant in the nervous system and, like brain NPF, promotes food intake, PYY is mainly produced by endocrine cells of the gut as a satiety factor. Our results indicate that *Drosophila* gut NPF, perhaps filling the role of mammalian gut PYY, acts to mediate sugar-specific satiety while promoting hunger for protein-rich yeast, illustrating a key hormonal mechanism that underlies selective hunger by which animals adjust their intake of specific nutrients.

Feeding decisions are based on internal state and exhibit sexual dimorphism. In *Drosophila*, males and females differ in their preference for and intake of dietary sugar and protein ^6^. Our findings define a complex interorgan communication system by which mating influences food choices in females. SP-induced female post-mating responses include decreased sugar intake and increased preference for protein-rich yeast food. Our results indicate that these responses to mating require midgut NPF signaling. When mated females consume dietary carbohydrates, NPF is released from the EECs and inhibits the AKH axis by directly suppressing AKH release from the APCs and by inhibiting release of midgut AstC, a factor that stimulates AKH secretion ^18^. Furthermore, NPF acts on the fat body to inhibit energy mobilization, thereby antagonizing AKH-mediated signaling in the adipose tissue. This inhibitory effect of NPF on AKH signaling reduces sugar consumption and promotes the intake of dietary yeast in mated females. While a number of studies have demonstrated that AKH is a regulator of metabolism [see ^2^], our findings uncover a key role of AKH in governing nutrient-specific feeding decisions. Recent work has revealed a sex-specific role of AKH, with lower activity in females underlying differences in male and female metabolism ^41^. Consistent with this notion, our results indicate that females use the midgut NPF system to suppress AKH signaling after mating to shift appetite towards protein-rich food. Furthermore, we recently showed that in mated females, midgut-derived AstC acts in a sex-specific manner through AKH to coordinate metabolism and food intake under nutritional stress ^18^. Our work here shows that NPF also works sex-specifically to sustain physiological requirements in females after mating by signaling from the gut to control AKH, suggesting that the gut-AKH axis occupies a central node in the hormonal circuit underlying sex-specific regulation of physiology.

How nutrient signals from the gut modulate feeding is key to understanding how nutritional needs are translated into specific feeding actions to maintain balance. We have identified a homeostatic circuit triggered by gut-derived NPF that limits sugar consumption and promotes the intake of protein-rich yeast. Similar mechanisms for sugar-induced satiety that drives protein hunger may also enable mammals to balance their intake of different nutrients with their metabolic needs. Elucidating how nutrient-responsive gut hormones such as NPF impact dietary choice is important to better understand hunger and cravings for specific nutrients that may ultimately lead to obesity.

## Methods

### *Drosophila* stocks and husbandry

Flies were reared on a standard cornmeal diet (82 g/L cornmeal, 60 g/L sucrose, 34 g/L yeast, 8 g/L agar, 4.8 mL/L propionic acid, and 1.6 g/L methyl-4-hydroxybenzoate) at 25 °C and 60% humidity with a 12-h light:12-h dark light cycle. Flies were transferred after eclosion to an adult-optimized diet (90 g/L sucrose, 80 g/L yeast, 10 g/L agar, 5 ml/L propionic acid, and 15 ml/L of 10% methyl-4-hydroxybenzoate in ethanol) ^48^ and aged to 4-7 days before experiments. Virgin female flies were collected within 3-5 hours of eclosion, whereas mated flies were sorted by sex a day before experiments. Genotypes that contained temperature-sensitive *Tubulin-GAL80*^*TS*^ were raised at 18 °C and kept on adult food for 3-4 days post-eclosion, after which they were incubated at 29 °C for five days to induce RNAi knockdown before experiments began. The animals were transferred to fresh food every third day. The following lines were obtained from the University of Indiana Bloomington *Drosophila* Stock center (BDSC): *AKH-GAL4* (#25684); *AstC::2A::GAL4* (#84595) ^24^; *UAS-mCD8::GFP* (#5137); *NPF::2A::GAL4* (#84671) ^24^; *elav-GAL4* (#458); *Cg-GAL4* (#7011); *da-GAL4* (#55850); CaLexA system (#66542: *LexAop-CD8::GFP::2A::CD8::GFP; UAS-LexA::VP16::NFAT, LexAop-CD2::GFP/TM6B, Tb*) ^30^; *UAS-TrpA1* (#26263 and a transgene insertion into *attP2*, a gift from C. Wegener); *UAS-NPF-RNAi*^*TRiP*^ (#27237); *UAS-NPFR-RNAi*^*TRiP*^ (#25939); *SP*^*0*^ mutant (#77892); *10XUAS-IVS-myr::tdTomato[su(Hw)attP8]* (#32223); and *Tub-GAL80*^*TS*^ (#7108). Other lines were obtained from the Vienna *Drosophila* Resource Center (VDRC): the control line *w*^*1118*^ (#60000, isogenic with the VDRC RNAi lines) as well as several *UAS-RNAi* lines including ones targeting *AKH* (#105063), *NPF* (*NPFi*^*KK*^, #108772, and *NPFi*^*sh*^, #330277), *NPFR* (*NPFRi*^*GD*^, #9605), *sut2* (#102028), and *Mondo* (#109821). *voilà-GAL4* ^49^ was a gift of Alessandro Scopelliti (University of Glasgow, UK). *R57C10-GAL80-6* ^50-55^ on the X chromosome was a kind gift of Ryusuke Niwa (University of Tsukuba). *AKH* mutant ^56^ and *NPFR::T2A::GAL4* ^32^ were gifts of Shu Kondo (Tokyo University of Science). *Df(3L)delta130* was a gift of Anne von Philipsborn (Aarhus University). For standardizing the genetic background and generating controls with proper genetic background, all GAL4 lines and GAL80 lines used this study were backcrossed for several generations to the same *w*^*1118*^ genetic background population before they were used in a final outcross with the genetic background of the RNAi lines and used as controls ^18^.

### Starvation-survival assay

Flies were transferred without anaesthesia to vials containing starvation medium (1% agar in water) and kept either at 29 °C (animals carrying *GAL80*^*TS*^) or at 25 °C (animals with constitutive GAL4 activity throughout development). Forty to 150 animals, at 10-15 flies per vial, were assayed for each genotype/sex. Dead animals were counted every 4-8 h. Statistical significance of survival differences was determined by using the Kaplan-Meier log-rank survival method in the Prism software package (version 9 or higher; GraphPad).

### Feeding Assays

Short-term food consumption was measured by using a spectrophotometric dye-feeding assay ^57,58^. During the morning meal (after lights-on), flies were transferred without anesthesia to adult-optimized food containing 0.5% erioglaucine dye (brilliant blue R, FD&C Blue No.1, Sigma-Aldrich, #861146) and allowed to feed for 30 minutes, if the flies had been 15 hours starved to stimulate food intake, or for 2-3 hours if not. Another set of flies were fed with undyed food to measure the baseline absorbance of fly lysates. For two-choice assays, the protocol of ^6^ was used with some modifications. Briefly, 25 flies were mildly anesthetized with CO_2_ before being placed in into a Petri dish (60 mm) with a checkerboard array of 20-µl patches of alternative diets containing either 100 g/L of sucrose, dyed red with 0.5% amaranth (Sigma #A1016), or 100 g/L yeast (dyed with 0.5% erioglaucine) and allowed to eat for 2 hours in the dark. For each genotype, 10-25 samples of 1-2 flies each were homogenized in 100 μL of phosphate buffer, pH 7.5, using a TissueLyser LT (Qiagen) bead mill with 5-mm stainless-steel beads (Qiagen, #69989). Homogenates were centrifuged at 16,000 *g* for 5 minutes, and 50 μL of each cleared supernatant was loaded into a 384-well plate. Sample absorbance was measured at 520 nm (amaranth) and at 629 nm (erioglaucine) on an Ensight multi-mode plate reader (PerkinElmer). Standard curves for erioglaucine and amaranth were used to convert absorbance values to food-consumption amounts.

Long-term food intake was monitored using the CAFÉ capillary-feeding assay ^23^. For consumption assays, assay tubes were constructed by inserting a 5-μL microcapillary (Hirschmann) through a hole in the lid of a 2-ml Eppendorf tube. The capillary was filled with a liquid sugar or yeast medium ^23^ containing 100 g/L sucrose or 100 g/L yeast extract, with 1 mL/L propionic acid and 1 g/L methyl-4-hydroxybenzoate preservatives, prior to the start of the experiment. For sugar preference assays, two capillaries were inserted into each tube, one filled with 10 g/L sucrose solution and the other filled with 100 g/L sucrose solution. Individual flies were briefly anesthetized on ice and placed into assay tubes, and the tubes were placed inside a moist chamber within a standard fly incubator. The level of the meniscus in each tube was measured at intervals. Tubes containing no flies were used as controls for evaporation; the amount of meniscus movement in these tubes was subtracted from the other measurements.

To monitor feeding behavior, interactions with food were measured over a 20–24 h period using the FLIC (Fly Liquid-Food Interaction Counter) assay ^21^. *Drosophila* Feeding Monitors (DFMs; Sable Systems, US) were installed in an incubator (25 °C, or 29 °C if *GAL80*^*TS*^ was present; 70% humidity, 12:12-hour light/dark cycle). Feeding wells were filled with a 10% sucrose solution, and individual flies were placed in each of the 12 chambers of the DFMs in the afternoon (after the morning meal) and left to acclimate for several hours, after which evening feeding data was recorded. The next morning, at lights on, fresh sugar solution was added to the DFMs, and morning meal data was recorded. In sugar-preference experiments, half of the DFM wells were filled with 1% sugar solution and the other half with 10% sugar solution. The data were recorded using the manufacturer’s software and analysed in *R Studio* using the published package, available at https://github.com/PletcherLab/FLIC_R_Code.

### Video recording of feeding behavior

Behavior chambers (40 mm in diameter) were coated with fluon on the top and sides to prevent flies’ walking on these surfaces. Fifteen-microliter patches of either 10% sucrose or 10% yeast (with no dyes) were placed in a circular pattern within the arena. Twelve 15-hour-starved animals per genotype were introduced into the chamber and were allowed to acclimate in darkness for a few minutes. The behavior chambers were placed on an infrared-light transilluminator viewed by a Basler camera, and half-hour videos were recorded at 15 Hz using the imaging setup described ^59^. Flies were tracked using the Ctrax software ^60^, and locomotion data were analyzed using custom MATLAB code.

### Immunohistochemistry and confocal imaging

Adult midguts, central nervous systems (CNSs), and fat bodies were dissected in cold PBS and fixed for 1 hour in 4% paraformaldehyde at room temperature with agitation. Anterior parts of guts, containing APCs (CCs) were first fixed for 30 min. Fixed tissues were quickly rinsed once with PBST (PBS with 0.1% Triton X-100, Merck #12298) and washed in PBST three times for fifteen minutes each. Washed tissues were incubated in blocking solution [PBST containing 5% normal goat serum (Sigma)] for 30 minutes at room temperature and incubated with primary antibodies diluted in blocking solution overnight (or 2 days for CNS samples) at 4 °C with gentle agitation. Primary antibody solution was removed, and the tissues were rinsed once and washed three times, 20 min each, with PBST. Tissues were incubated with secondary antibodies diluted in PBST overnight at 4 °C, washed three times with PBST, and mounted in Vectashield mounting medium containing DAPI (Vector Laboratories, #H-1200) on slides treated with poly-L-lysine (Sigma, #P8920). Tissues were scanned on a Zeiss LSM-900 confocal microscope using a 20× air objective using the Zen software package. Image analysis was performed by using the open-source program ImageJ ^61^. For quantification of NPF, AstC, AKH, and GFP (CaLexA reporter) staining intensity, samples to be compared were stained simultaneously using the same reagent preparations and imaged with the same settings. Stacks were Z-projected using the sum method. For gut analyses, a custom MATLAB script was used to determine the number of stained cells and their intensity. To subtract local background, the brightness of the area surrounding each identified EEC was measured and subtracted from each measured cell intensity. We integrated a *UAS-tdTomato* transgene into the CaLexA system to normalize calcium-dependent GFP fluorescence with GAL4 activity intensity. Antibodies used included a rabbit antibody against the processed AKH peptide ^36^, a kind gift of Jae Park, U. Tennessee, 1:500; rabbit anti-NPF (Ray BioTech, #RB-19-0001-20), 1:500; rabbit anti-AstC ^62^, kindly given by Jan Veenstra, U. Bordeaux, and Meet Zandawala, Brown University, 1:500; mouse anti-GFP (ThermoFisher #A11120), 1:500; rat anti-mCherry (used against tdTomato; ThermoFisher, #M11217), 1:2000; mouse anti-Prospero (University of Iowa Developmental Studies Hybridoma Bank, #MR1A), 1:20; Alexa Fluor 488-conjugated goat anti-mouse (ThermoFisher, #A32723), 1:500; Alexa Fluor 555-conjugated goat anti-rabbit (ThermoFisher #A32732), 1:500; Alexa Fluor 555-conjugated goat anti-rat (ThermoFisher, #A21434), 1:500; and Alexa Fluor 405-conjugated goat anti-rabbit (ThermoFisher, #31556), 1:500.

### Injection experiments

Synthetic amidated NPF peptide (SNSRPPRKNDVNTMADAYKFLQDLDTYYGDRARVRFamide) was a kind gift of Frank Hauser (U. Copenhagen). Peptide was dissolved at 25 μM in a synthetic hemolymph-like buffer ^63^ containing 5 mM glucose, 5 mM trehalose, and 110 mM sucrose (inert osmolyte). Flies of (1) *Tub-GAL80*^*TS*^; *voilà*>*NPF-RNAi*^*KK*^ and (2) *Tub-GAL80*^*TS*^; *voilà*>*NPF-RNAi*^*sh*^, along with corresponding controls *Tub-GAL80*^*TS*^; *voilà*>*+*, were reared at 18 °C as described above. After 4 days at 29 °C to induce RNAi knockdown, flies were starved on 1% agar for 15 hours. Starved flies were anaesthetized on ice, and 50 nL of hemolymph-like solution with or without NPF was injected into each animal at the lateral mid-thorax ventral to the wings using a Nanoject II injector (Drummond Scientific, PA, USA). Assuming each injected animal contained one microliter of hemolymph, the final NPF concentration in the injected animals was increased to 1.25 μM, a level that should strongly activate NPFR [IC_50_ ∼60 nM ^64^]. Animals were allowed to recover for 30 minutes at 29 °C before use in dye-feeding assays as described above.

### Metabolite measurements

Triglyceride and glycogen levels were measured using established protocols ^22,65^. For each genotype, 10 batches of 3 flies each were homogenized in PBS containing 0.05% Tween-20 (Sigma #1379) in a TissueLyser LT (Qiagen) bead mill with 5-mm stainless-steel beads. Glycogen was measured by hydrolyzing glycogen into glucose by using amyloglucosidase (Sigma, #A7420) followed by colorimetric glucose measurement (Sigma, #GAGO20). Triacylglyceride levels were assayed by cleaving their ester bonds using Triglyceride Reagent (Sigma, #T2449) to obtain free glycerol, the level of which was colorimetrically measured using the Free Glycerol reagent (Sigma, #F6428). For determination of circulating glucose concentration, hemolymph was extracted as described previously (Tennessen et al., 2014), and glucose was measured using the colorimetric assay (Sigma, #GAGO20). Each sample’s absorbance at 540 nm was measured in a 384-well plate using an Ensight multimode plate reader (PerkinElmer) and converted to metabolite concentrations using glycerol and glucose standard curves. Measurements are reported on a per-fly basis.

### Transcript measurement using qPCR

Six tissue replicates (each containing five CNSs, five midguts, or five CC-containing anterior parts of guts) per condition or genotype were homogenized in 2-ml Eppendorf tubes containing lysis buffer with 1% beta-mercaptoethanol using a TissueLyser LT bead mill (Qiagen) and 5-mm stainless steel beads (Qiagen #69989). RNA purification was performed using the NucleoSpin RNA kit (Macherey-Nagel, #740955) according to the manufacturer’s instructions. Complementary DNA was synthesized using the High-Capacity cDNA Synthesis kit (Applied Biosystems, #4368814). QPCR was done using RealQ Plus 2x Master Mix Green (Ampliqon, #A324402) on a QuantStudio 5 (Applied Biosystems) machine. Results were normalized against the housekeeping gene *Rp49* using the delta-delta-Ct method. The oligos used are listed in Table S1.

### Statistics

All the statistics were computed using the Prism analysis package (GraphPad). Starvation-survival curves were analyzed using Kaplan-Meier log-rank tests or Gehan-Breslow-Wilcoxon test. Other data were assessed for normality before comparisons were performed. For normally distributed data, pairwise comparisons were made using two-tailed Student’s t-tests, and multiple samples were compared using one-way ANOVA with post-hoc multiple-comparisons tests. Other data were compared using two-tailed Mann-Whitney U tests or one-way Kruskal-Wallis ANOVA. To assess interactions between experimental variables, two-way ANOVA was conducted with Bonferroni’s multiple comparisons as a post-hoc test. Plots show the mean with standard error of the mean (SEM). Significance is shown as: ns, *p*≥0.05; *, *p<*0.05; **, *p<*0.01; ***, *p<*0.001.

## Acknowledgements

Anti-Akh was a kind gift of Jae Park (University of Tennessee). Anti-AstC was a generous gift of Jan Veenstra (University of Bordeaux) and Meet Zandawala (Brown University). *R57C10-GAL80* was kindly given by Ryusuke Niwa (University of Tsukuba). *AKH*-mutant and *NPFR::2A::GAL4* flies were kindly provided by Shu Kondo (Tokyo University of Science). We thank the Vienna *Drosophila* Resource Center and the University of Indiana Bloomington *Drosophila* Stock Center for fly lines, and we also thank the University of Iowa Developmental Studies Hybridoma bank for providing anti-Prospero. This work was supported by Novo Nordisk Foundation grant NNF19OC0054632 and Lundbeck Foundation grant 2019-772 to KR. TK and KVH were supported by funding from the Villum Foundation (15365) and Danish Council for Independent Research Natural Sciences (9064-00009B) to KVH.

## Extended figures

**Fig. S1.**
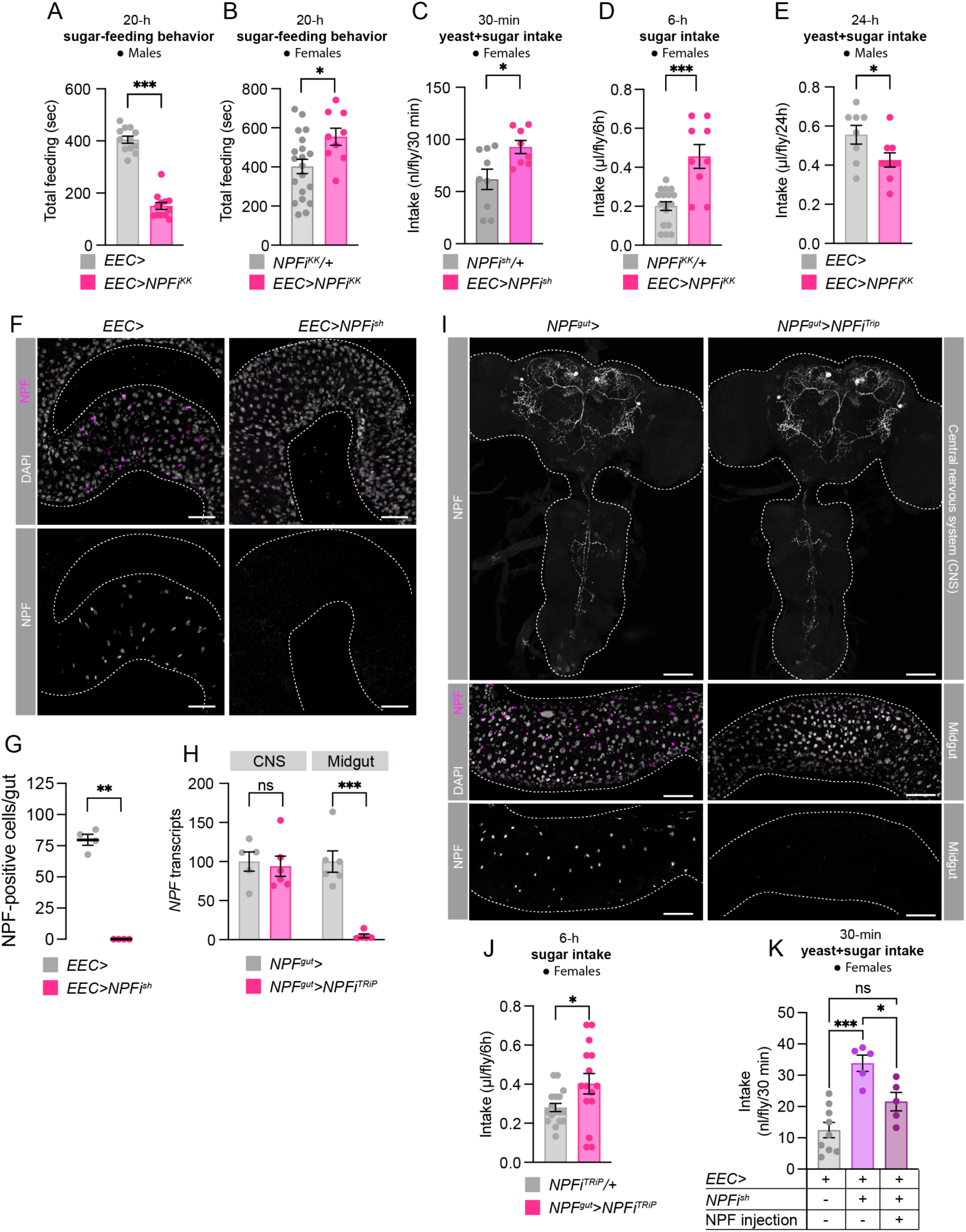
*NPF* knockdown efficiently depletes NPF specifically in the midgut EECs and affects food intake. (A) Total feeding time of males using FLIC on sugar food. Student’s t-test; n=12 *EEC*>, n =12 *EEC*>*NPFi*^*KK*^. (B) Time spent feeding on 10% sugar food by mated females measured using FLIC over 24 hours. Student’s t-test; n=20 *NPFi*^*KK*^*/+*, n=9 *EEC*>*NPFi*^*KK*^. (C) Consumption of sugar+yeast food (9% sugar + 8% yeast) by 15-hour-starved mated females over 30 min determined by dye assay. Student’s t-test; n=9 *NPFi*^*sh*^*/+* and n=8 *EEC*>*NPFi*^*sh*^. (D) Consumption of 10% sugar by 15-hour-starved mated females measured over 6 hours by CAFÉ assay. Mann-Whitney U test; n=17 *NPFi*^*KK*^*/+* and n=9 *EEC*>*NPFi*^*KK*^. (E) Consumption of sugar+yeast food (5% sugar + 5% yeast extract) by males over 24 hours measured by CAFÉ assay (Student’s t-test; n=8 *EEC*>, n=9 *EEC*>*NPFi*^*KK*^). (F and G) NPF immunostaining of midguts from mated females with knockdown of *NPF* demonstrates that *EEC*>-driven RNAi using *NPFi*^*sh*^ completely abolishes NPF signal in the midgut, quantified in G (Mann-Whitney U test; n=4 guts). Scale bars 50 µm. (H) Knockdown using the *R57C10-GAL80, NPF*> (*NPF*^*gut*^>) driver with *NPFi*^*TRiP*^ strongly reduces midgut *NPF* transcripts in mated females without affecting expression in the CNS (brain and ventral nerve cord). One-way ANOVA with Kruskal-Wallis test (n=5 *NPF*^*gut*^> CNS, n=6 *NPF*^*gut*^>*NPFi*^*TRiP*^ CNS, n=6 *NPF*^*gut*^> midgut, n=5 *NPF*^*gut*^>*NPFi*^*TRiP*^ midgut, independent biological replicates from tissues pooled from 6 animals). (I) NPF immunostaining of CNS and midguts from mated females with knockdown of *NPF* show that *NPF*^*gut*^>-driven *NPFi*^*TRiP*^ completely abolishes NPF signal in the midgut but does not affect CNS levels, quantified in Fig. 1H. Scale bars 50 µm. (J) Consumption of 10% sugar by 15-hour-starved mated females over 6 hours measured by CAFÉ assay. Student’s t-test; n=17 *NPFi*^*TRiP*^*/+*, n=15 *NPF*^*gut*^>*NPFi*^*TRiP*^. (K) Injection of NPF peptide into the hemolymph restores normal feeding without affecting hemolymph glucose levels. Consumption of sugar+yeast food over 30 minutes in 15-hour-starved mated females. One-way ANOVA with Tukey’s test; n=9 *EEC*>, n=5 *EEC*>*NPFi*^*sh*^, n=5 *EEC*>*NPFi*^*sh*^*+*NPF peptide. All females were mated. Error bars indicate standard error of the mean (SEM). ns, non-significant; *, *p*<0.05; **, *p*<0.01; and ***, *p*<0.001.

**Fig. S2.**
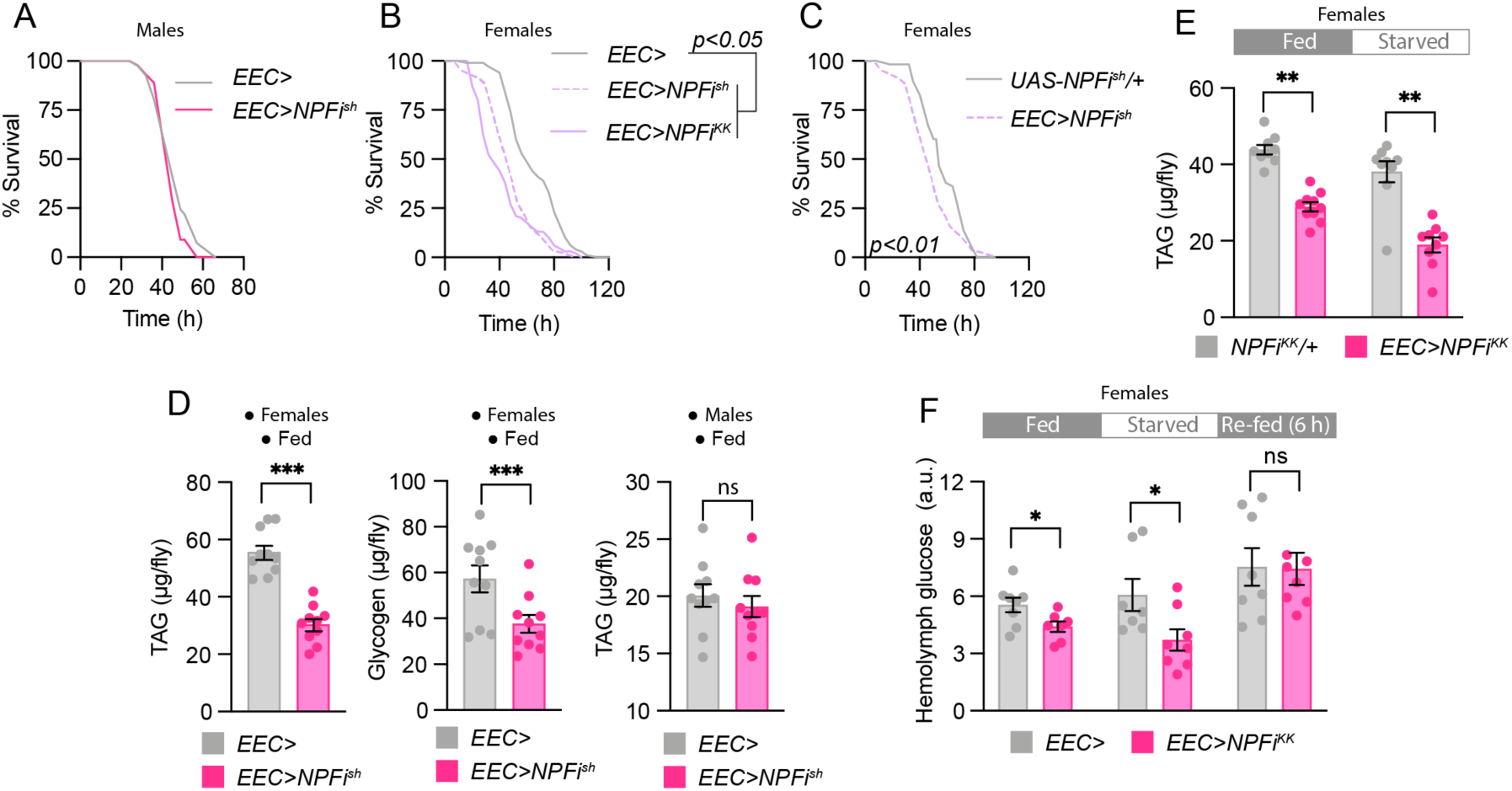
Enteroendocrine NPF regulates resistance to nutritional stress and systemic metabolism. (A-C) Survival during starvation compared using Gehan-Breslow-Wilcoxon test (A, n=41 *EEC*> males, n=45 *EEC*>*NPFi*^*sh*^ males; B, n=130 *EEC*>, n=122 *EEC*>*NPFi*^*sh*^, n=94 *EEC*>*NPFi*^*KK*^; C, n=50 *UAS-NPFi*^*sh*^*/+*, n=122 *EEC*>*NPFi*^*sh*^). (D) TAG and glycogen levels in flies in the fed condition. Mann-Whitney U tests (n=10 independent biological replicates with 3 animals per sample). (E) TAG levels in mated females in the fed condition and following 24-hour starvation. Mann-Whitney U tests (n=9 fed *NPFi*^*KK*^*/+*, n=10 fed *EEC*>*NPFi*^*KK*^, n=9 starved *NPFi*^*KK*^*/+*, n=9 starved *EEC*>*NPFi*^*KK*^). (F) Circulating hemolymph sugar levels in mated females in the fed condition, following 24 h starvation, and after 6 h of re-feeding from the 24-h-starved condition. Student’s t-tests (n=8 fed *EEC*>, n=7 fed *EEC*>*NPFi*^*KK*^, n=7 starved *EEC*>, n=8 starved *EEC*>*NPFi*^*KK*^, n=8 re-fed *EEC*>, n=7 re-fed *EEC*>*NPFi*^*KK*^ independent biological replicates with 3 animals per sample). All females were mated. Error bars indicate standard error of the mean (SEM). ns, non-significant; *, *p*<0.05; **, *p*<0.01; and ***, *p*<0.001.

**Fig. S3.**
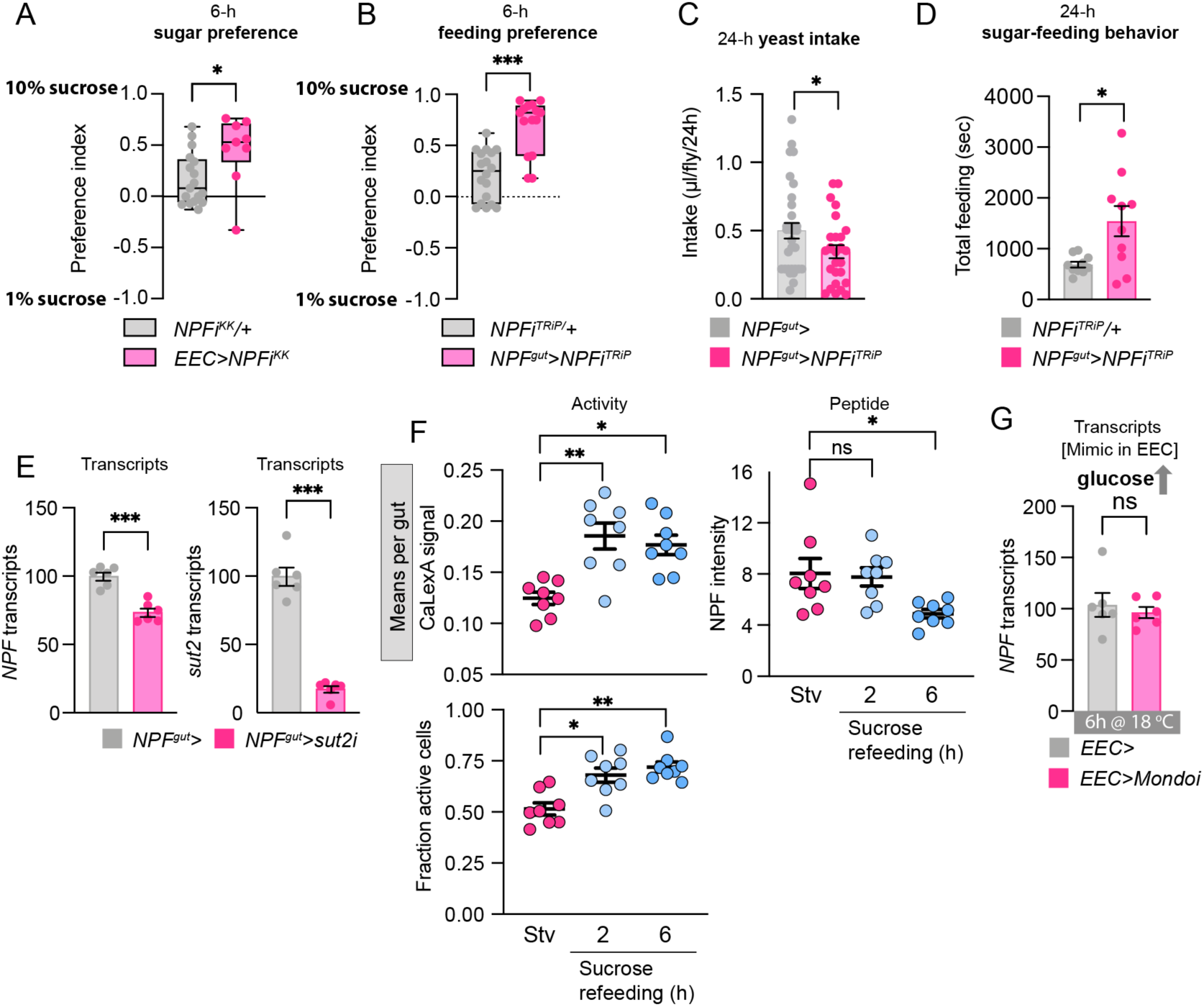
Gut loss of *NPF* affects feeding and is influenced by sugar sensing. (A-B) Consumption preference of mated females for 1% and 10% sugar after 15 hours’ starvation, measured over 6 hours by CAFÉ assay. Mann-Whitney U tests (n=17 *NPFi*^*KK*^*/+*, n=9 *EEC*>*NPFi*^*KK*^, n=18 *NPFi*^*TRiP*^*/+*, n=15 *NPF*^*gut*^>*NPFi*^*TRiP*^). (C) Consumption of 10% yeast by 15-h starved mated females over 24 hours determined by CAFE. Mann-Whitney U test (n=27 *NPF*^*gut*^>, n=26 *NPF*^*gut*^>*NPFi*^*TRiP*^). (D) Feeding time on 10% sucrose of 15-h starved mated females measured by FLIC over 24 hours. Student’s t-test (n=10 *NPFi*^*TRiP*^*/+*, n=10 *NPF*^*gut*^>*NPFi*^*TRiP*^). (E) Transcript levels of *NPF* (left) and *sut2* (right) in midguts from fed mated females. Student’s t-test (n=6 replicates containing five tissues each). (F) NPF intensity and NPF-cell activity (CaLexA) in 24-h starved mated females and 2-h- and 6-h-sugar-re-fed mated females measured by calcium-LexA reporter system, aggregated on a per-gut basis. One-way ANOVA with Kruskal-Wallis test. Eight guts per condition. (G) Transcript levels of *NPF* in midguts from fed mated females. Five midguts in each replicate. Student’s t-test (n=5 *EEC*>, n=6 *EEC*>*Mondoi*). All animals were mated females. Error bars indicate standard error of the mean (SEM). ns, non-significant; *, *p*<0.05; **, *p*<0.01; and ***, *p*<0.001.

**Fig. S4.**
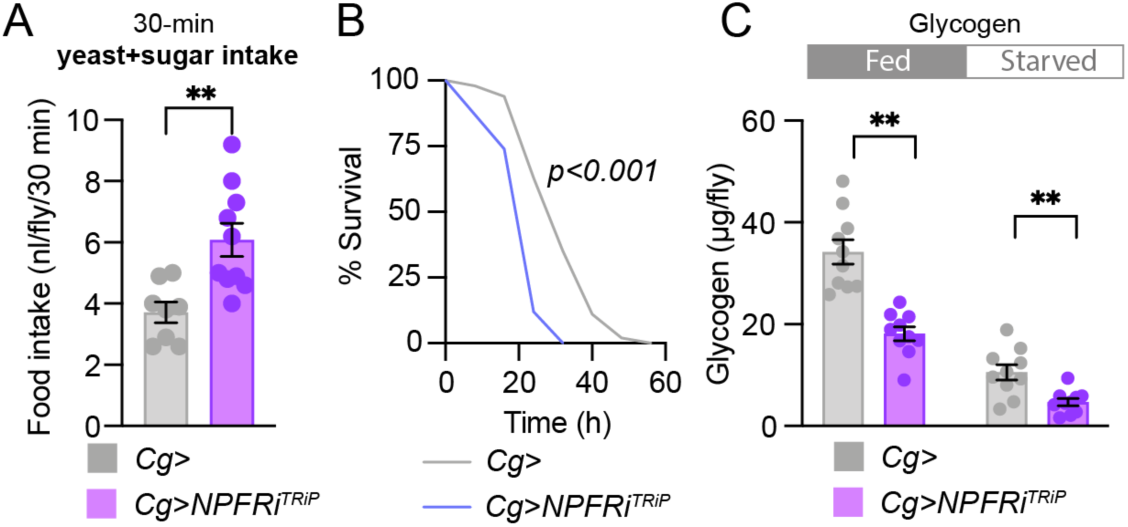
Knockdown of *NPFR* in the fat leads to increased food intake after starvation, starvation survival sensitivity, and lower glycogen stores. (A) Food intake by 15-h-starved mated females over 30 min, measured by dye assay (9% sugar and 8% yeast). Student’s t-test (n=8 *Cg*>, n =10 *Cg*>*NPFRi*^*TRiP*^). (B) Survival during starvation of mated females compared using Gehan-Breslow-Wilcoxon test (n=100 *Cg*>, n =50 *Cg*>*NPFRi*^*TRiP*^). (C) Glycogen levels in fed and 15-h starved mated females. One-way ANOVA with Dunnett’s post-hoc test, all n=10. All animals were mated females. Error bars indicate standard error of the mean (SEM). ns, non-significant; *, *p*<0.05; **, *p*<0.01; and ***, *p*<0.001.

**Fig. S5.**
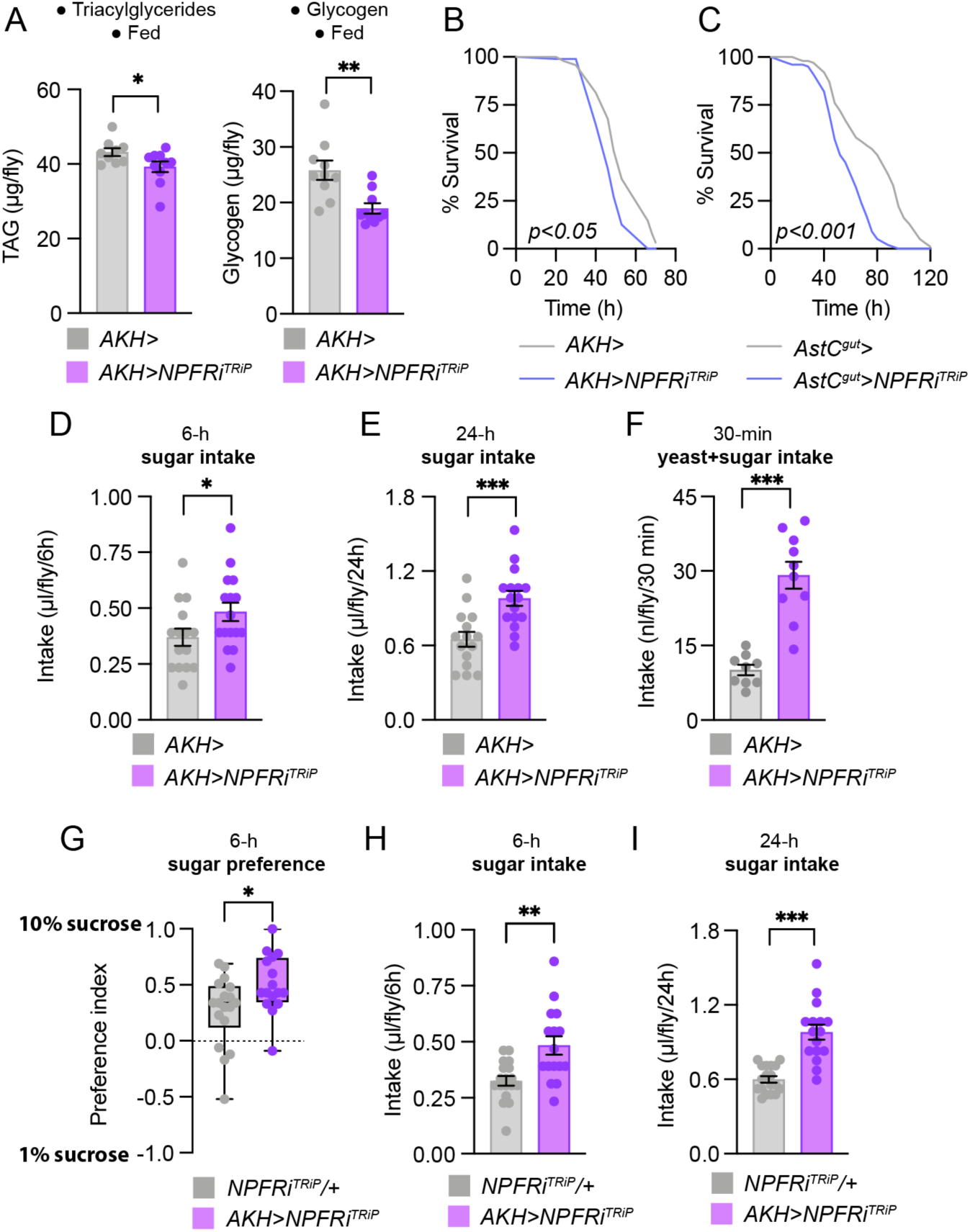
Knockdown of *NPFR* in the AKH-producing cells leads to starvation survival sensitivity, reduced energy stores, and increased preference for and intake of dietary sugar. (A) Metabolite levels in fed mated females. TAG: Student’s t-test (n=9 *AKH*>, n=10 *AKH*>*NPFRi*^*TRiP*^). Glycogen: Mann-Whitney U test (n=10 *AKH*>, n=10 *AKH*>*NPFRi*^*TRiP*^). (B and C) Survival during starvation of mated females compared using Gehan-Breslow-Wilcoxon test (n=96 *AKH*>, n=96 *AKH*>*NPFRi*^*TRiP*^, n=153 *AstC*^*gut*^>, n=198 *AstC*^*gut*^>*NPFRi*^*TRiP*^). (D and E) Consumption of 10% sucrose by 15-h-starved mated females over 6 hours and 24 hours measured by CAFÉ assay. Mann-Whitney U test (n=15 *AKH*>, n=16 *AKH*>*NPFRi*^*TRiP*^). (F) Food intake (9% sugar and 8% yeast) by 15-hour-starved mated females measured over 30 min by dye assay. Student’s t-test (n=9 *AKH*>, n=10 *AKH*>*NPFRi*^*TRiP*^). (G) Consumption preference of 15-hour-starved mated females for 1% and 10% sucrose measured by CAFÉ assay over 6 hours. Student’s t-test; n=18 *NPFRi*^*TRiP*^*/+*, n=16 *AKH*>*NPFRi*^*TRiP*^. (H and I) Consumption of 10% sucrose by 15-h-starved mated females measured over 6 hours and 24 hours by CAFÉ assay. Mann-Whitney U tests (n=18 *NPFRi*^*TRiP*^*/+*, n=16 *AKH*>*NPFRi*^*TRiP*^). All animals were mated females. Error bars indicate standard error of the mean (SEM). ns, non-significant; *, *p*<0.05; **, *p*<0.01; and ***, *p*<0.001.

**Fig. S6.**
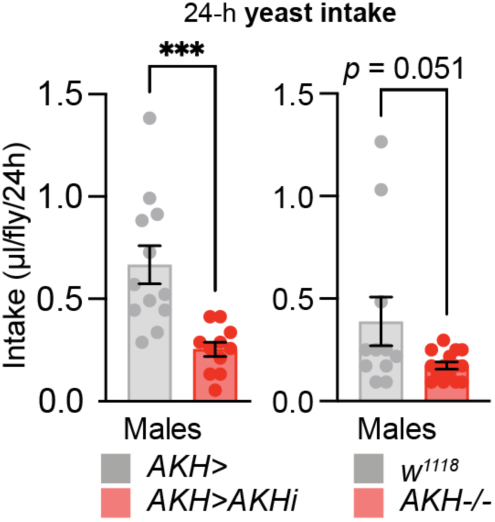
Loss of AKH leads to reduced yeast intake in males, indicating sexual dimorphism in AKH effects on feeding preferences. Yeast intake of 15-hour-starved males measured over 24 hours by CAFÉ assay with 10% yeast extract medium. Left panel, *AKH* knockdown; right panel, *AKH* mutant. Student’s t-test; n=12 *AKH*>, n=11 *AKH*>*AKHi*^*KK*^, n=11 *w*^*1118*^, n=15 *AKH*^-/-^. Error bars indicate standard error of the mean (SEM). ***, *p*<0.001.

**Table S1:**
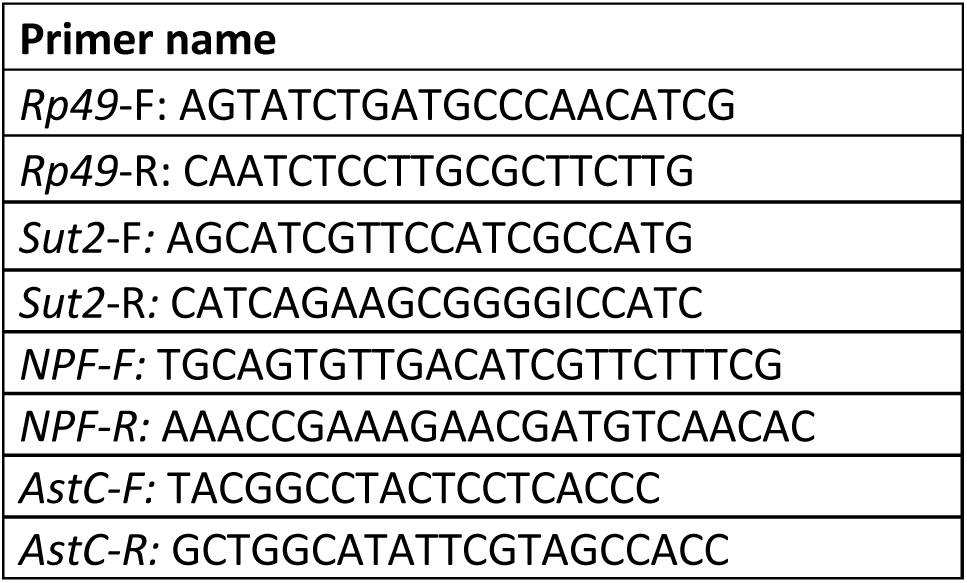
Oligo sequences used for qPCR.

